# Astrocyte-induced internal state transitions reshape brainwide sensory, integrative, and motor computations

**DOI:** 10.64898/2026.02.05.704034

**Authors:** Jing-Xuan Lim, Ziqiang Wei, Sujatha Narayan, Yan Zhang, Jeremy P. Hasseman, Ilya Kolb, Jihong Zheng, Alireza Sheikhattar, Xuelong Mi, Wei Zheng, Xueying Yang, Mariam Beriashvili, Greg Fleishman, Caroline L. Wee, Chris de Zeeuw, Guoqiang Yu, Behtash Babadi, Mikail Rubinov, Loren L. Looger, Dwight E. Bergles, James E. Fitzgerald, Misha B. Ahrens

**Affiliations:** Janelia Research Campus, Howard Hughes Medical Institute, Ashburn, VA, USA; The Solomon H. Snyder Department of Neuroscience, Johns Hopkins University School of Medicine, Baltimore, MD, USA; Institute of Molecular and Cell Biology (IMCB), Agency for Science, Technology and Research (A*STAR), Singapore; Allen Institute for Brain Science, Seattle, WA, USA; GENIE Project Team, Janelia Research Campus, Howard Hughes Medical Institute, Ashburn, VA, USA; Department of Electrical and Computer Engineering, University of Maryland, USA; Bradley Department of Electrical and Computer Engineering, Virginia Tech, Blacksburg, VA, USA; Neuroscience Institute, New York University Langone Medical Center, New York, NY, USA; Department of Neuroscience, Erasmus MC, Rotterdam, The Netherlands; Netherlands Institute for Neuroscience, Royal Academy of Sciences, Amsterdam, The Netherlands; Department of Biomedical Engineering, Vanderbilt University, Nashville, TN, USA; Department of Computer Science, Vanderbilt University, Nashville, TN, USA; Department of Neurosciences, School of Medicine, University of California, San Diego, CA, USA; Kavli Neuroscience Discovery Institute, Johns Hopkins University, Baltimore, MD, USA; Department of Neurobiology, Northwestern University, Evanston, IL, USA; NSF-Simons National Institute for Theory and Mathematics in Biology, Chicago, IL, USA

**Author notes:** Equal contribution (Jing-Xuan Lim, Ziqiang Wei, James E. Fitzgerald, Misha B. Ahrens).

**Keywords:** Internal state, astrocytes, evidence accumulation, behavioral state, network dynamics, sensory processing, sensorimotor transformation, motor preparation, motor control

## Abstract

Animals rapidly adapt to changing circumstances by shifting how they perceive, integrate, and act. Such flexibility is often attributed to transitions between internal states that exert widespread influence across the brain. Yet the mechanisms that drive state transitions and how they reconfigure brainwide computation remain unclear. Larval zebrafish, when actions are rendered futile by decoupling visual flow feedback from swimming in virtual reality, enter a temporary passive, energy-preserving state. In this state, astrocyte calcium levels are elevated, and swim reinitiation requires greater accumulated visual motion. Using whole-brain, cellular-resolution activity imaging, we observed widespread circuit alterations underlying this disengaged state: neuronal visual responses weakened, visual motion integration over time became dramatically leakier, motor inhibition increased, and motor preparation slowed, together suppressing conversion of sensory evidence into action. Astrocyte calcium rose during futile swimming, tracked the emergence and resolution of these brainwide changes, and was both necessary and sufficient to drive them. Thus, astrocytes orchestrate internal states that profoundly reshape neural computations, most powerfully at intermediate integrative processing stages, to meet changing demands.

**Highlights:**

- Internal state change alters brainwide neuronal processing at every stage of the sensorimotor transformation
- Effects are most powerful at integrative stages through stimulus memory collapse
- As state resolves, amplification of sensory representations synergizes with reduced motor inhibition for action reinitiation
- Astrocyte activity drives these brainwide adaptive shifts in neuronal dynamics

## Introduction

The constantly changing environment and body of an animal requires it to act and respond to incoming stimuli in flexible ways. Such behavioral flexibility can be induced via ‘internal states’ such as alertness, fear, attentional states, and hunger—variables that are not directly observable but that exert broad influence over perception, cognition, and behavior,^1^ and which typically persist beyond individual stimuli or actions.^2^ In principle, an internal state might be mediated by a localized change in a brain circuit, while on the other extreme, it might be mediated by widespread, coordinated changes in sensory, memory, decision, and motor planning circuits.^3^ At the neural level, internal states are known to shape network activity and sensory responses.^4–8^ State changes can be induced by neuromodulator release, allowing anatomically stable circuitry to exhibit variations in dynamics to support behavioral adjustments on relatively short time scales.^9–19^ Despite observations of certain anatomically and functionally localized correspondences between internal states and circuit activity, it is virtually unknown how global computations—spanning perception, information accumulation, motor preparation, and action—are jointly modulated, in part due to the difficulty of measuring neural activity with cellular resolution across all brain regions in behaving animals, and the challenges of dissociating internal states from ongoing behavior. Furthermore, the causal network- and molecular mechanisms by which internal states arise from recent experience and exert their modulatory effects on brainwide circuit dynamics remain largely unknown.

To address these questions at the whole-brain level, we imaged cellular activity throughout the entire brain of larval zebrafish, a vertebrate that experiences a variety of internal states.^20–30^ We examined neuronal and astroglial activity patterns during futility-induced passivity, a disengaged state induced when swim actions consistently fail to produce their expected outcomes, resulting in temporary behavioral passivity.^27^ This behavioral switch is the result of noradrenergic signaling for action failures and the integration of this futility signal by astrocytes.^27^ We developed a fast calcium sensor, jGCaMP8p, whose rapid kinetics enabled clear separation of individual sensory responses from temporally integrated activity. We found that during the futility-induced passive state there were coordinated changes in the dynamics of vast numbers of neurons across the whole brain, spanning activity related to sensation, integration, and action. Remarkably, this modulation of circuit activity, despite being global and coordinated across brain regions, was highly specific to the computation that neurons were involved in. The most dramatically impacted computation was visual motion integration over time—which normally induces compensatory swimming to stabilize fish location—a dynamic process that was suppressed many-fold in the futility-induced disengaged state, primarily through a change in the leakiness of integration, i.e., a shortening of the memory time constant.^31,32^ Molecular gain- and loss-of-function experiments revealed that astrocyte calcium increases were both necessary and sufficient to induce these large-scale changes in circuit dynamics. These results indicate that astrocyte activation globally modulates neuronal computations across the entire brain during internal state transitions and show that astrocytes orchestrate broad, adaptive shifts in circuit function.

## Results

### Futility-induced modulation of behavioral responses in a sensory accumulation task

We employed a futility-induced passivity behavioral assay to elicit internal state changes and compared an altered, disengaged state that arises after experiencing behavioral futility (Fig. 1A, bottom) with a normal, engaged state (Fig. 1A, top). This assay is based on the tendency of larval zebrafish to swim against water flow and in the direction of optic flow to stabilize their location in space, a response also present in virtual-reality (VR) environments.^33–36^ Mathematically, this can be represented as *v_stim_=v_flow_−m×G_ms_*, with *v_stim_* indicating the velocity of the projected visual scene, *v_flow_* the velocity of the flow, *G_ms_* the motosensory gain (the speed of visual feedback per unit motor output), and *m* the motor vigor, which was recorded electrophysiologically from the tail motor nerve in a fictive swimming preparation (Fig. 1B; Methods). Instead of a closed-loop VR where *G_ms_>0* (Fig. 1C, left), setting *G_ms_=0* creates an open-loop environment in which the fish is unable to control its position as the visual scene continues to move forward with *v_stim_=v_flow_*, i.e. swimming is futile (Fig. 1C, right). After multiple swim attempts, fish tend to ‘give up’ and remain behaviorally passive for several tens of seconds (futility-induced passivity) before recovering to an active state.^27,37^

**Figure 1.**
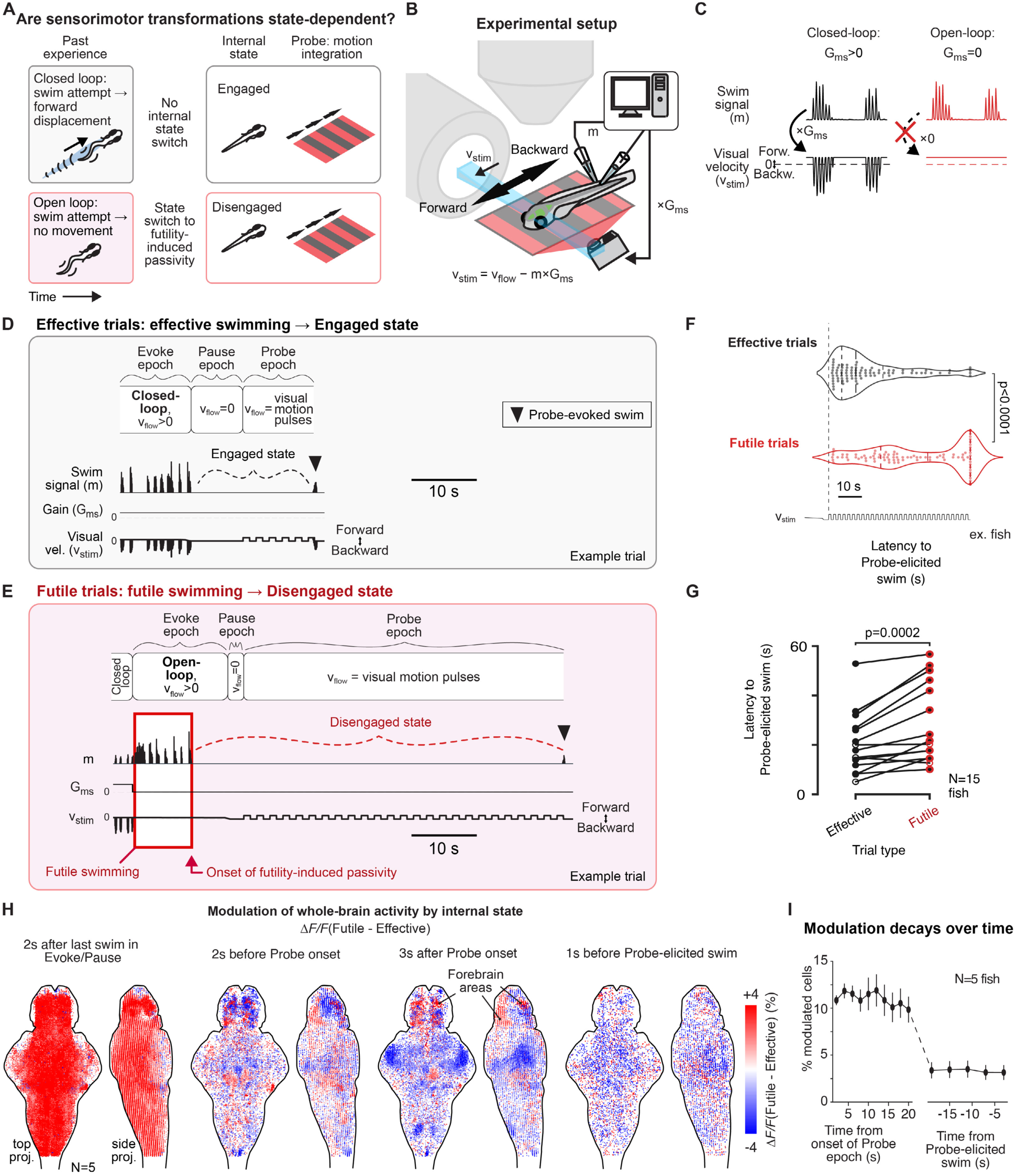
Effects of futility-induced passivity on behavior and brain activity in a visual accumulation task. **(A)** Conceptual paradigm. Effective swimming in a closed-loop environment (top) maintains an “Engaged” internal state. Futile swims in an open-loop environment will result in a “Disengaged” state and behavioral passivity. This study compares brainwide sensorimotor processing under these two behaviorally quiescent yet distinguishable internal states. **(B)** Experimental setup. Fish were placed in a virtual reality (VR) environment where action-outcomes were controlled to evoke Engaged or Disengaged states. Velocity of the projected visual scene is determined by *v_stim_*=*v_flow_−m×G_ms_* with *v_stim_* the stimulus speed, *v_flow_* the simulated flow, *m* fictive motor output, and *G_ms_* the multiplicative gain factor between motor output and stimulus motion. **(C)** Illustration of closed-loop VR (*G_ms_>0*; left; black lines) where swims lead to visual feedback simulating forward body displacement and open-loop VR (*G_ms_=0*; right; red lines) where swim attempts had no effect on the stimulus. **(D-E)** Behavioral assay to study internal state-dependent sensorimotor processing consisting initially of closed-loop swimming to ensure an Engaged internal state. In the Evoke epoch, either an Engaged state was maintained (Effective trials, **D**) or a Disengaged state was induced (Futile trials, **E**). The Pause epoch allowed direct motor and visual effects on brain activity to decay. In the Probe epoch, fish on both trial types were exposed to slow pulsatile forward visual motion; fish remained quiescent for some time in both Engaged (D) or Disengaged (E) states but tended to respond later in the Disengaged state. **(F-G)** Fish exhibited delayed responses to forward visual motion pulses during the Disengaged state. **(F)**. Top, violin plots of the time from Probe onset to the Probe-evoked swim (i.e. latency to Probe-elicited swim), with the quartiles indicated as vertical dashed lines. Dot, recovery time of a single trial. Probe epochs were truncated at 60 s if fish did not swim; rank sum test. **(G)** Statistics of average latencies to Probe-elicited swim across fish. Dots, average latencies to Probe-elicited swims across trials in individual fish. Solid dots, fish with significant differences of recovery times in Effective and Futile trials (p<0.05; sum rank test, 12 fish), empty dots, fish with insignificant differences of recovery times (p>0.05; sum rank test, 3 fish); N=15 fish; paired signed rank test on all 15 fish. **(H)** Snapshots of state-dependent dynamics across fish (N=5) at (i) 2 s after the offset of the last swim in either the Evoke or Pause epoch, (ii) 2 s before the onset of the Probe epoch, (iii) 3 s after the onset of the Probe epoch, to (iv) 1 s before the fish swims in the Probe epoch. Blue-red colormap, *ΔF/F_Futile_−ΔF/F_Effective_<0*, in blue; *ΔF/F_Futile_−ΔF/F_Effective_>0*, in red. Data from individual fish were registered to a reference brain which was then applied as a mask. Activity maps show differences in the Probe epoch determined by internal state, and that these differences became smaller over time, and mostly equalized by the time of behavioral response, which occurs on average later on Futile trials. **(I)** Percentage of state-dependent neurons decreases over time, shown aligned to Probe onset (left) or Probe-evoked swim onset (right). A neuron is defined to be state-dependent if its response is significantly different during Effective and Futile trials at a given time (p<0.05; rank sum test). See also Figure S1.

To examine the influence of internal states on subsequent neural processing of incoming stimuli and their transformation into actions, we designed a paradigm in which fish underwent such changes in internal states and then, during behavioral quiescence, performed a sensory temporal integration task.^38–42^ Trial types alternated between “Effective” trials and “Futile” trials designed to evoke different internal states prior to the sensory accumulation task.

On Effective trials (Fig. 1D; Methods), fish first swam in a closed-loop (*G_ms_>0*) environment during the Evoke epoch, designed to maintain a normal internal state in which fish engage with visual motion stimuli—the “Engaged” state. Next, in the Pause epoch where *v_flow_=0*, fish typically stopped swimming for some time.^43,44^ Then, during the Probe epoch, a series of slow, one-second forward-moving visual motion stimuli were shown, alternating with one second periods of no motion. Fish were behaviorally quiescent while they integrated the visual motion pulses and eventually responded to it by swimming (Fig. 1D).

Futile trials unfolded similarly (Fig. 1E) and started in a closed-loop environment but had one crucial difference: in the Evoke epoch, the environment switched to open-loop where swimming was futile and no longer triggered visual feedback (*G_ms_=0*) which evoked a futility-induced “Disengaged” state, where fish ‘gave up’ and stopped swimming for a period and disengaged from interacting with the visual motion stimuli.^27^ This automatically-detected transition (Methods) triggered the Pause epoch, followed by the previously described Probe epoch in which fish integrated a series of visual motion pulses before responding.

Importantly, during the Probe epoch of both Effective and Futile trials, animals exhibited similar behavioral quiescence but occupied different internal states that could then be related to distinct brain dynamics. We used this paradigm to define the effects of these internal state differences on sensory integration and behavior.

### Internal state gates behavioral responses to visual motion

On Futile trials, fish in the Disengaged state responded with a longer latency to the series of forward motion pulses in the Probe epoch compared to the Engaged state (Effective trials, 21.8 ± 3.3 s; Futile trials, 31.5 ± 4.2 s; mean ± SD; N=15 fish; Fig. 1F-G and Fig. S1A). Thus, internal state modulated behavioral responses during the sensory integration task.

To rule out the possibility that behavioral differences arose simply from differences in visual history, we introduced open-loop replay trials, during which the *v_stim_* time series from the Evoke epoch of the last Effective trial was replayed in open-loop replay Futile trials. These trials also led to futility-induced passivity, followed by delayed recovery times (Fig. S1A). Thus, the experience of behavioral futility, specifically, triggers the change in internal state.

### Internal state is reflected in brainwide neuronal dynamics

To define the effects of internal state on neural dynamics throughout the brain, we performed whole-brain light-sheet imaging^36^ during the task. To dissociate fast sensory from integrative components of the sensorimotor transformation, we developed a new, ultra-fast calcium indicator, jGCaMP8p that, when expressed in neurons under the pan-neuronal *elavl3* promoter, decayed within one second, which is the time between visual motion pulses in the Probe epoch, and could therefore be used to disambiguate neural representations of visual velocity and pulse-by-pulse neural integration of visual velocity over time (Fig. S1B-D).^45^

During the Evoke epoch, overall activity was higher on Futile trials, likely due to increased motor vigor (Fig. 1H, first panel), but this activity difference had declined in most brain regions before the Probe epoch (Fig. 1H, second panel). During the Probe epoch, neural activity was generally weaker on Futile trials (Fig. 1H, third panel). Just before the probe-evoked swim (i.e. generally at a later time in Futile trials), activity across the brain was similar between Futile and Effective trials (Fig. 1H, fourth panel), suggesting resolution of the Disengaged internal state. The number of neurons with activity differences (p<0.05, rank sum test of ΔF/F in Futile versus Effective trials) changed from 12 ± 1% at the start of the Probe epoch to 3 ± 1% just before the probe-evoked swim (Fig. 1I).

### Distributed circuits transform sensory input into motor output

To characterize the effects of internal state on neural dynamics from perception to action, we used a simplified model as a guide to analyze whole-brain data (Fig. 2A1) based on (i) encoding of sensory input, (ii) integration of that information over time, (iii) preparation for the motor response, and (iv) generation of motor output.

**Figure 2.**
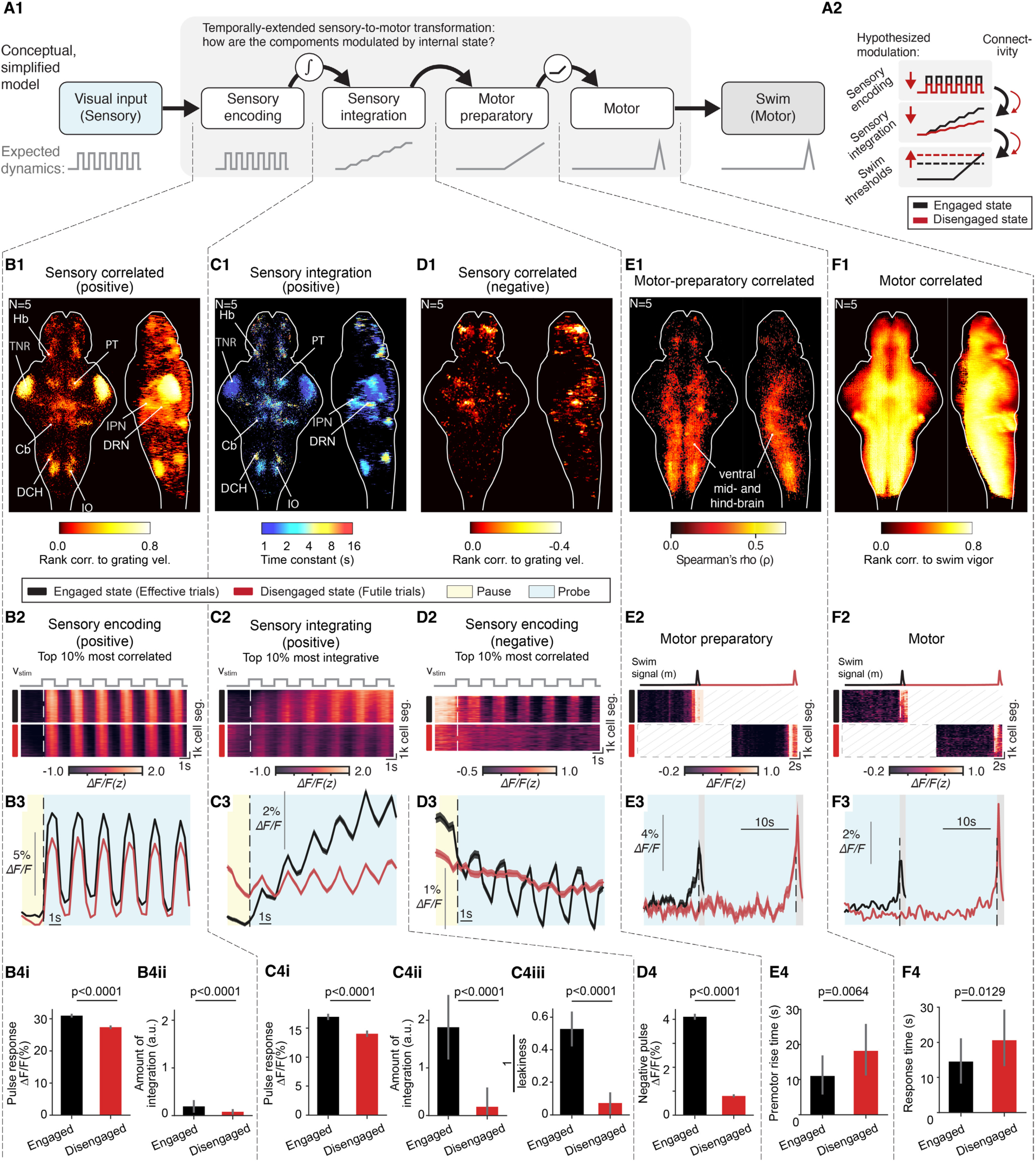
Brainwide state-dependent dynamics at all stages of sensorimotor processing. **(A) (A1)** Schematic of a simple feedforward conceptual model for sensorimotor processing. **(A2)** Subset of hypotheses for the ways that internal state may impact the different processing stages. **(B)** State-dependent sensory encoding in positive sensory-responding cells. **(B1)** Brain map of rank correlation between neuronal responses and time series of forward motion pulses (⍴>0, p<0.05); N=5 fish. **(B2)** Activity of the top 10% highest rank correlation positive sensory-responding cells; top, forward motion pulses; middle, Effective trials (black bar); bottom, Futile trials (red bar); N=5 fish. **(B3)** Average dynamics across neurons in (B2); Effective trials, black; Futile trials, red; shaded area, SD across neurons; yellow, Pause epoch; blue, Probe epoch. **(B4i)** Pulse responses (area under curve of a single pulse starting from pulse onset). **(B4ii)** Bar plots of pulse integration (area under curve of the baseline dynamics in the first 6 pulses, i.e. the neural dynamics after removing single pulse response from **(B4i)**). **(C)** State-dependent sensory integration. Positive sensory-responding cells are shown. **(C1)** Brain map of decay time constants of neuronal sensory responses for neurons in (B). **(C2)** Activity of the top 10% positive sensory-responding cells with longest decay time constants; same convention as (B2). **(C3)** Average dynamics across neurons in (C2); same convention as (B3). Integration on Futile trials (red) almost vanishes. **(C4)** Statistics of pulse responses **(C4i)**, pulse integration **(C4ii)**, and integration leakiness (**C4iii**, depicted as 1/leakiness; Methods) across neurons in (C2); the same convention as (B4). **(D)** State-dependent sensory integration in negative sensory-responding cells. **(D1)** Brain map of rank correlations between neuronal response and motion pulses (⍴<0, p<0.05); N=5 fish. **(D2)** Activity pattern of top 10% negative sensory-responding cells with the highest negative rank correlations; same convention as (B2). **(D3)** Average dynamics across neurons in (D2); same convention as (B3). **(D4)** Bar plot of negative pulse responses (negative area under curve of the dynamics in the first 6 pulses from the onset of the first pulse); the same convention as (B4). **(E)** State-dependent motor preparatory activity. **(E1)** Brain map of rank correlation between motor preparatory neuronal dynamics and latencies to the Probe-evoked swims (p<0.05, ⍴>0; top 5% cells, neural dynamics during Engaged state; Methods). **(E2)** Activity of motor preparatory cells in example fish. **(E3)** Average dynamics across neurons in (E2) with probe-evoked swim timing difference presenting the trial average. **(E4)** Bar plot of average times when motor preparatory cell activity starts to rise; p=0.0064, two-way ANOVA, N=5 fish; the same convention as (B4). **(F)** State-dependent motor-related activity. **(F1)** Brain map of positive Spearman’s rank correlations between neuronal responses and swim vigor during *G_ms_=0* (motor-related cells, ⍴>0, p<0.05); N=5 fish. **(F2)** Activity pattern of motor-related cells; the same convention as (E2). **(F3)** Average dynamics across neurons in (F2) with probe-evoked swim timing difference presented as average difference of swim timing in Futile vs. Effective trials; conventions as (E3). **(F4)** Bar plot of average probe-evoked swim times read out from motor cells; p=0.0129, two-way ANOVA (Methods), N=5 fish; same conventions as (E4). See also Fig. 4E, negative brain map. See also Figure S2 for more detail about motor preparatory activity.

Sensory-encoding neurons were determined by the significance (p<0.05) of their Spearman’s correlation between each neuron’s activity and *v_stim_* during Probe. Positive sensory-responding neurons were present in multiple brain areas including the tectal neuropil region (TNR), habenula (Hb), superior raphe (DRN), interpeduncular nucleus (IPN), cerebellum (Cb), dorsocaudal hindbrain (DCH) including Spatial-Location encoding Medulla Oblongata neurons (SLO-MO), inferior olive (IO), pretectum (PT) and other regions (Fig. 2B1).^41,46,47^

Sensory-integration neurons were characterized via the integrative time constant, *τ*, measured by fitting the neural response of sensory-encoding neurons to the integrated series of forward motion pulses during the Probe epoch as *ΔF/F(t) ∼ ∫v_stim_(t-t’)e^-t’/τ^dt’*, where *τ=∞* implies perfect retention of total displacement (perfect integration of velocity) and small *τ* implies responses to velocity that are gradually forgotten (leaky integration). The integrative time constant *τ* is shown, for each neuron in the dataset, in Fig. 2C1, and shows dependence on brain region.

Negatively-responding sensory and integrative neurons (Fig 2D1) were present in smaller numbers (positively-responding cells: 11 ± 3% of all neurons; negatively-responding cells: 3 ± 2% of all neurons; N=5 fish) and showed mild integrative properties (Fig. 2D3).

Putative “motor preparatory” signals with ramping activity several seconds before swimming (Fig. S2A-B; Methods) were distributed throughout the brain, especially in the ventral mid- and hindbrain (Fig. 2E1). Activity here remained low but started rising ∼3 seconds before fish swam (Fig. 2E3; Fig. S2A,B).

Finally, motor-related signals were found to be widely distributed across brain areas (Fig. 2F1).^46,48^ Some of this activity likely directly drove swimming, while other neuronal populations likely received motor efference copies from spinal circuits or motor command neurons in the brain.^49^

### The Disengaged state modulates all stages of the sensation-to-action process but most profoundly impacts temporal stimulus integration

In principle, the delayed responses during the Disengaged state could arise from changes anywhere along the sensorimotor pathway (Fig. 2A2). To define the effects of internal state on the neural dynamics of neurons along the sensation-to-action model, we first focused on the Probe epoch, where fish received the same stream of visual input and were behaviorally quiescent regardless of internal state.

At the sensory encoding level, we observed an overall dampening effect of futility-induced passivity on sensory responses, where positive sensory-responding neurons responded less to visual motion pulses in the Disengaged state compared to the Engaged state (17 ± 1% lower response amplitude in the Disengaged state; N=5 fish; Methods) (Fig. 2B2, 2B3, 2B4i). This shows that the visual system still encoded the pulses in the Disengaged state, but less strongly than before.

At the temporal integration level, positive stimulus-integrating neurons showed a dramatic reduction in activity of over 7-fold in the Disengaged state (87 ± 3% lower, Fig. 2C2-C3). This reduction was already present in neurons with stronger sensory encoding relative to integration (70 ± 3%, Fig. 2B4ii), and this effect was further enhanced in neurons with stronger sensory integration, which showed an over 9-fold reduction in integration (90 ± 5%, Fig. 2C4ii). Since the immediate response to visual motion pulses were only mildly suppressed in this population (Fig. 2Ci) compared to the dramatic suppression in overall integration (Fig. 2C4ii), the reduction in integration is unlikely inherited from the reduction in sensory responses (e.g. Fig. 2B4i), and more likely to be due to changing integration properties. Indeed, we observed an increase in integration leakiness (Fig. 2C4iii), i.e. a shortening of the integration time constant leading to faster forgetting (average integration time constant in the Engaged state: 11.2 ± 4.7 s; in the Disengaged state: 5.9 ± 4.3 s). This suggests that changes in integration time constants play an important role in altered information retention and processing during different internal states.^31^

The effect of internal state on sensory-negative cells was also dramatic, showing a greater than 5-fold reduction (82 ± 4%) of this downward sensory response during the Disengaged state (Fig. 2D2-D4).

Finally, activity of motor preparatory as well as the motor-related populations was delayed in the Disengaged state, consistent with behavior (Fig. 2E2-E4, 2F2-F4). Additionally, weak visual responses were observed within the motor preparatory population of Fig. 2E1, which were also state-modulated (Fig. S2C,D). Overall, it took longer for motor preparatory cells to initiate ramp-up dynamics during Futile trials, possibly due to weaker activation resulting from leakier integration in upstream regions.

These results indicate that the futility-induced Disengaged internal state impinges on all stages of the sensorimotor transformation, but—through a profound increase in leakiness, reduced information retention, and a sevenfold overall dampening—affects the temporal integration stage most strongly.

### Region specific modulation of neural circuit dynamics

To investigate the roles of individual brain areas in state-dependent sensorimotor processing, we performed clustering of neural responses and focused on sensory and integrative responses (Fig. 3A; more details in Fig. S3).^50^ Cells encoding or integrating visual motion pulses were modulated by different degrees and for different durations, illustrated by three example areas (Fig. 3B). We examined the depth of modulation at the single-cell level using the discriminability index (d’, Methods), confirming that most of these cells were suppressed in the Disengaged relative to the Engaged state (Fig. 3C) and almost none exhibited the opposite property (Fig. S4A; 3±2%). Depth of modulation (Fig. 3D) correlated with the percentage of integrative cells in each region (Fig. 3E; Spearman’s r=0.68, p=0.037).

**Figure 3.**
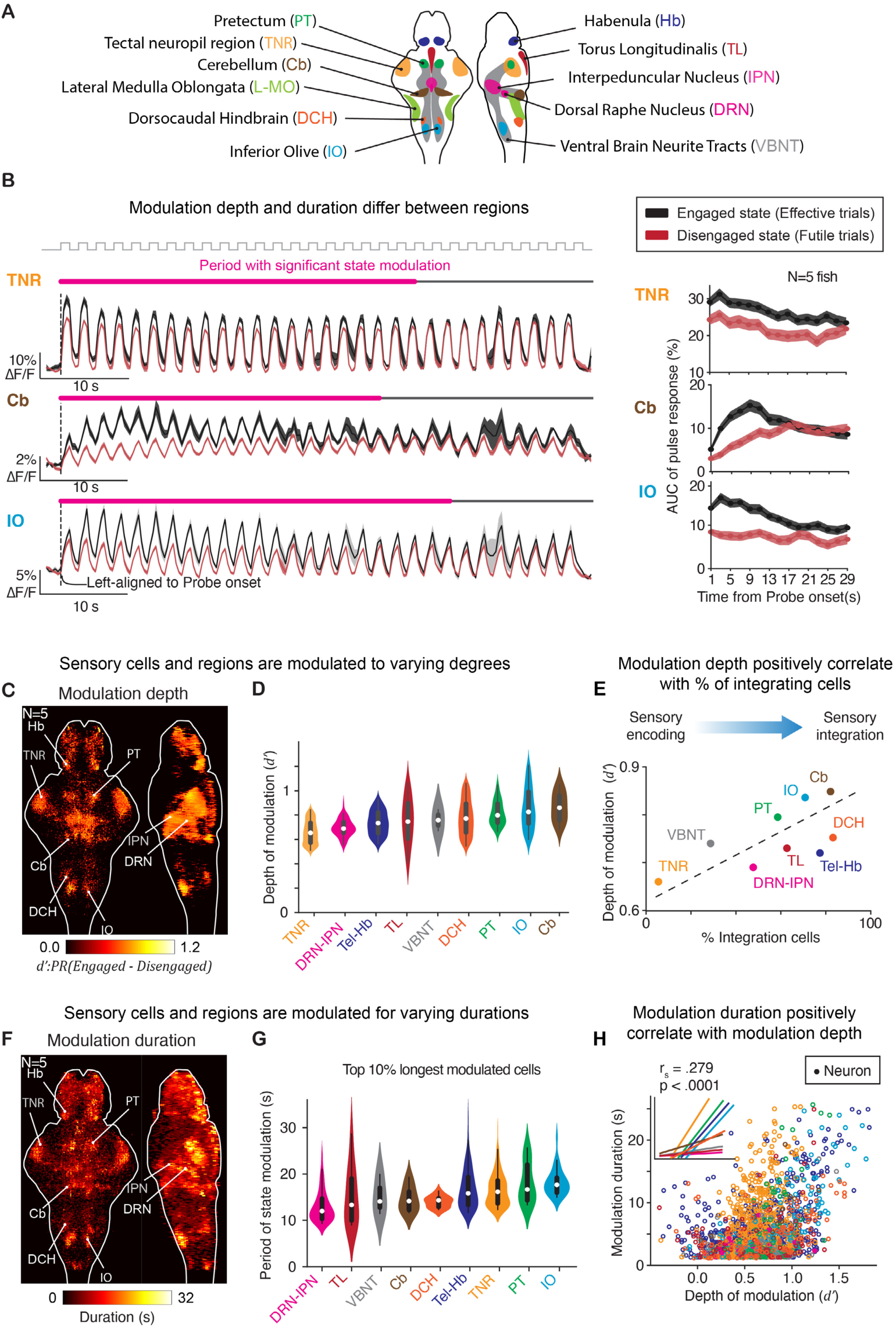
Internal states modulate distributed brain dynamics to different extents and for different durations. **(A)** Schematized, labeled example brain regions showing correlates to sensorimotor processing, derived from a spatiotemporal clustering algorithm. **(B)** Effect of internal state on sensory processing decreases over time in Probe epoch. Left, average activity of three example regions. Magenta line, period of time when activity shows modulation by internal state (rank sum test; p<0.05); gray line, period of time when state modulation is insignificant. Right, amplitudes of response to individual pulse over time. Black, Engaged; red, Disengaged. Shaded area, SEM; N=5 fish. Neural dynamics centered at Evoke onset across fish. Measured as decrease of pulse responses for the first 5 forward motion pulses from passive relative to Engaged states, cells were state-modulated by 18.47±8.82% (mean±SD; N=5 fish) in TN, 52.56±5.41% in Cb, and 46.72±7.59% in IO. **(C-E)** Depths of state modulation vary across brain regions and are correlated to sensory integration. Depth is quantified by the discriminability index (d’: PREngaged−PRDisengaged) of neural responses to the visual motion (PR for “pulse response”) at the early phase of the Probe epoch. **(C)** Brain map of positive d’ across fish (N=5 fish). **(D)** Distribution of d’ in sensory processing brain regions. Violin plots, ordered by their average d’; empty circles, averages of d’; box-and-whisker plots, 1^st^ and 3^rd^ quartiles of the distributions. **(E)** Depth of modulation is positively correlated with the proportion of sensory cells with long decays (>4 s) in each region; dashed line, linear regression. **(F-H)** Duration of state modulation varies across brain regions but increases with depth of modulation within a given brain region. **(F)** Brain map of modulation duration across fish (N=5 fish). **(G)** Distribution of durations in sensory processing brain regions, same convention as **(D)**. **(H)** Within regions, duration of modulation is correlated with depth of modulation. Inner plot, linear regressions of duration versus depth of modulation on single neurons. See also Figures S3, S4 and S5.

We also determined the period of modulation by internal state at the single-cell level, i.e. the time period before neural responses to motion pulses in Futile trials became indistinguishable from responses in Effective trials (Fig. 3F). Across brain regions, the top 10% longest modulated cells were modulated for >10 s in the Disengaged state (Fig. 3G), with consistent effects visible when inspecting all cells (Fig. S4B). Across single cells within regions, larger depth of modulation was associated with longer duration of modulation (Fig. 3H). This suggests that a global causal variable determines internal state, and that the return to normal brain dynamics unfolds nonhomogeneously across brain regions.

A population within the DCH region, SLO-MO neurons, known to integrate and memorize visual flow over long time periods, showed higher Probe-unrelated activity in Futile trials (Fig. S5), reflecting the additional ≥5 seconds of forward visual motion exposure during the Evoke epoch, consistent with their role in sensory integration.^41^

### Progressive amplification of visual responses predicts behavior

Animals eventually returned to a behaviorally active state and resumed swim attempts, at which point brain activity between the two trial types was mostly equalized (Fig. 1H and 4A, right). To compare activity across trial types shortly after the internal state switch and at the varying times of motor recovery, we plotted visual responses across the entire Probe epoch, aligned on the left to Probe onset and aligned on the right to the visual pulse that triggered motor recovery. In the middle, we plotted time-warped data to account for varying trial lengths. We found that, eventually, the visual responses in Effective and Futile trials became indistinguishable, but this happened on average at a later time point on Futile trials (Fig. 4A).

**Figure 4.**
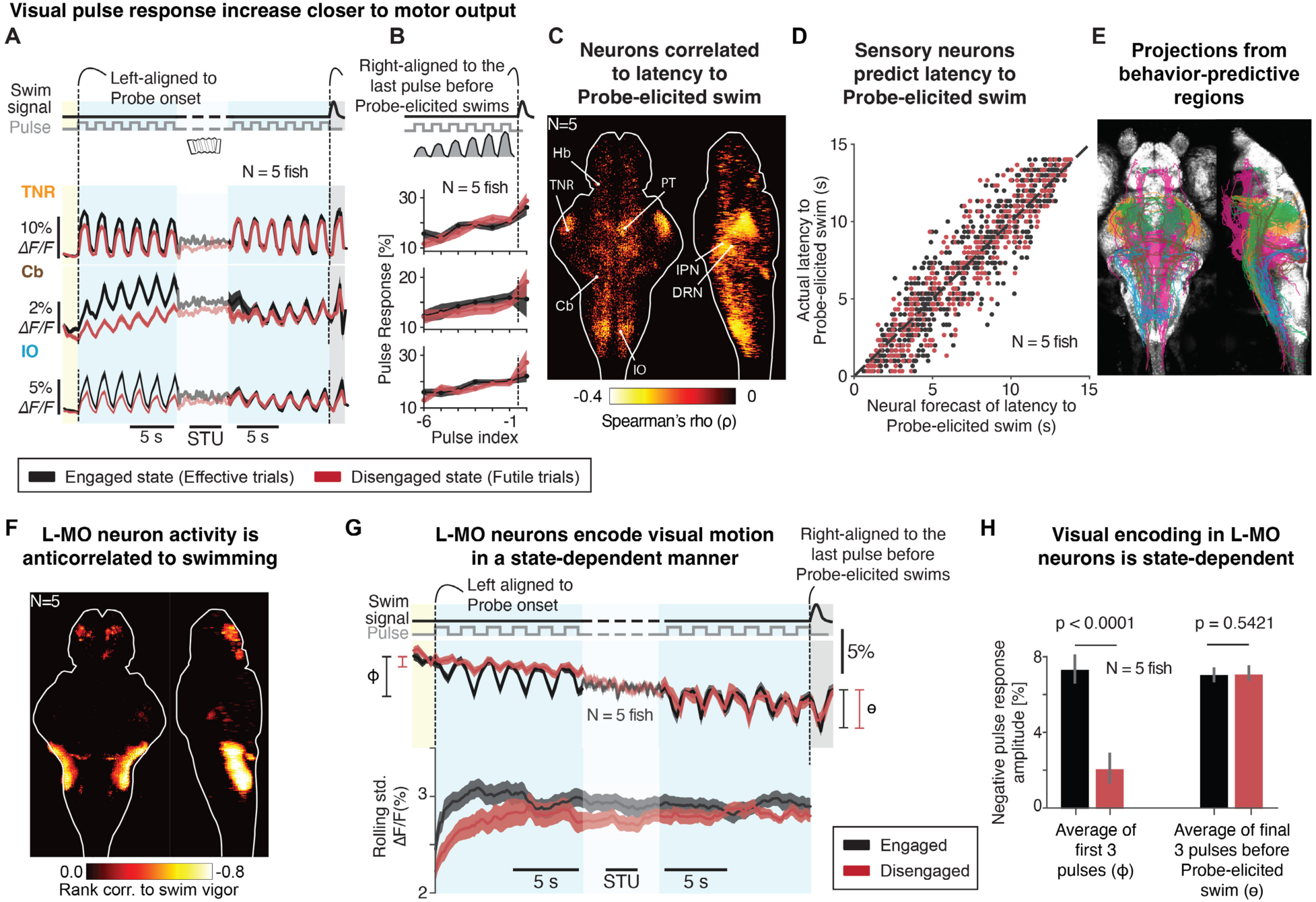
Dynamics of sensory regions during the state-recovery process. **(A-B)** Neural responses on Futile trials eventually resembled those on Effective trials at a later time, followed by progressive amplification of responses preceding probe-evoked swims. **(A)** Yellow, Pause epoch; blue, Probe epoch. Left, trials aligned to Probe onset; right, aligned to the last pulse before probe-evoked swims; middle, time-warped dynamics in between (STU: scaled time unit). **(B)**, neural responses to motion pulses (area under curve per pulse) were amplified progressively before Probe-evoked swims. Shaded area, SEM across fish; N=5 fish. Pulse index: pulse number relative to Probe-elicited swim. TNR, tectal neuropil region; Cb, cerebellum; IO, inferior olive. Neural activity was centered at the Evoke onset across fish. **(C)** Brain map of behavior-predictive sensory regions whose neuronal responses to a randomly-selected pulse in Probe is predictive of the latency from that pulse to the probe-evoked swim; Spearman’s rank correlation; ⍴<0, p<0.05. Behavior-predictive sensory regions include TNR, PT, Cb, DRN, IPN, and IO. **(D)** Latency to probe-evoked swims decoded from neural responses to visual motion (Methods). Y-axis, actual latency from a given pulse to the probe-evoked swim; x-axis, predicted latency from neural dynamics across all behavior-predictive sensory regions. Dots, single trials; dashed line, unity (x=y). Effective trials, black; Futile trials, red; N=5 fish. **(E)** Putative synaptic recipients of the behavior-predictive sensory regions whose neuronal cell bodies reside in TNR (yellow), PT (green), Cb (brown), DRN or IPN (magenta), and IO (cyan) from the mapzebrain atlas (https://mapzebrain.org/). **(F)** Brain map of the negative rank correlation between neuronal responses and swim vigors during *G_ms_=0* (motor-negative cells, ⍴<0, p<0.05); N=5 fish. The dominant region in this brain map is located in the lateral medulla oblongata (L-MO). **(G)** State-dependent negative visual response in L-MO. Negative visual response was reduced in the early phase of Probe epoch in Futile trials. Top, average neural dynamics, centered to activity at Evoke onset across fish; bottom, rolling standard deviation in a 4-second time window. Effective trials, black; Futile trials, red; shaded area, SEM across fish; N=5 fish. Left blue, aligned to Probe onset; right blue, aligned to the last pulse before the probe-evoked swim; middle, time-warped activity in between (STU: scaled time unit). **(H)** Bar plots of the state-dependent visual response in L-MO neurons. Left, early pulse response in Probe (as marked in left side of (F)), p<0.0001, paired signed rank test across neurons; right, late pulse response (as marked in right side of (F)), p<0.5421, paired signed rank test across neurons. Effective trials, black; Futile trials, red; error bar, SD across neurons, N=5 fish. See also Figure S6 for more analyses of L-MO neuronal dynamics.

In both trial types, visual response amplitude increased in the lead-up to behavioral recovery (Fig. 4A,B).^51^ This suggests that sensory responsiveness of individual neurons correlated with motor preparedness, indicating a sequence in which a recovery from a Disengaged to an Engaged state occurred first, followed by a visually-induced behavioral preparatory process. Furthermore, we found high correlations between visual response amplitudes in the lead-up to swimming and probe-elicited swim times in cells in regions across the brain such as TNR, PT, Cb, DRN, IPN and IO (Fig. 4C). A decoder of all sensory-responding cells (Methods) could accurately predict behavioral recovery time (Fig. 4D), suggesting that the progressive increase in visual response amplitude was part of the mechanism underlying visually-driven swim initiation.

Using mapzebrain (https://mapzebrain.org/), we visualized the morphology of neurons in these areas, revealing that projection patterns were broadly consistent with the functional data (Fig. 4E, cf. Fig. 2E1).^52,53^

Thus, brainwide activity underwent consistent changes in the lead-up to behavioral responses, suggesting widespread engagement and modulation of diverse neuronal networks involved in the timing and execution of visually-driven behavioral responses.

### Internal state controls sensory gating in a motor preparation-related population

In addition to increasing sensory responsiveness (Fig. 4B) and ramping motor preparatory activity (Fig. 2E1), we found state-dependent modulation of more complex activity related to swim generation in the lateral medulla oblongata (L-MO). L-MO neuron activity was anticorrelated with swim vigor (Fig. 4F), consistent with its known activation by astroglia (Fig. S6A) and role in suppressing swim motor output during futility-induced passivity ^27^. Activity was also spontaneously oscillatory with a consistent phase relationship to swim initiation, suggesting an additional role in the timing of routine swim bouts (Fig. S6B-D). This region responded to forward motion pulses, albeit unreliably, with a decrease in activity (Fig. S6E; note that L-MO is absent from d’ and sensory responding maps due to its stochastic sensory response), which, due to L-MO’s inhibitory effect on swimming, suggests that this sensory-driven disinhibition of motor output contributes to the behavioral response to visual motion. Remarkably, in the Disengaged state, these negative visual responses were almost completely absent (Fig. 4G,H), i.e. “gated-off”, after which they started to appear over the course of the Probe epoch as the “gate” gradually opened, and were robustly present at the moment of full recovery (Fig. 4G,H). This finding suggests that L-MO neurons play a role both in generating precisely timed motor output by cyclically suppressing and releasing motor circuits, and in internal state-dependent control of sensory-driven behavior.

Together, these findings indicate that in the Disengaged state, sensory encoding is mildly reduced, integration becomes much leakier and provides weaker drive to motor-preparatory circuits, while motor suppression from L-MO is enhanced and its sensory inhibition gated off (Fig. S6F). As the Disengaged state recedes, sensory encoding and integration recover, motor-inhibitory circuits quieten, and sensory-driven motor disinhibition is restored. Once normalized, sensory integration and increasing sensory sensitivity can again drive motor output. Thus, the internal state manifests as modulation of circuit dynamics spanning sensation to action, producing broad yet highly specific effects on subcircuits.

### Brainwide neuromodulatory and astroglial correlates to the Disengaged state

Consistent with astrocytes driving futility-induced passivity behavior based on noradrenergic signaling for futile swimming, astroglial calcium rose during the experience of behavioral futility (Fig. 5A-B), remained elevated during the Pause and Probe epochs (Fig. 5C,D), and had returned to baseline when fish recovered back to the Engaged state (Fig. 5E; neuromodulator dynamics shown in Fig. S7).^27,28,37,54–63^ We therefore hypothesized that astrocytes drove the observed alterations in brainwide neural dynamics during the Disengaged state.

**Figure 5.**
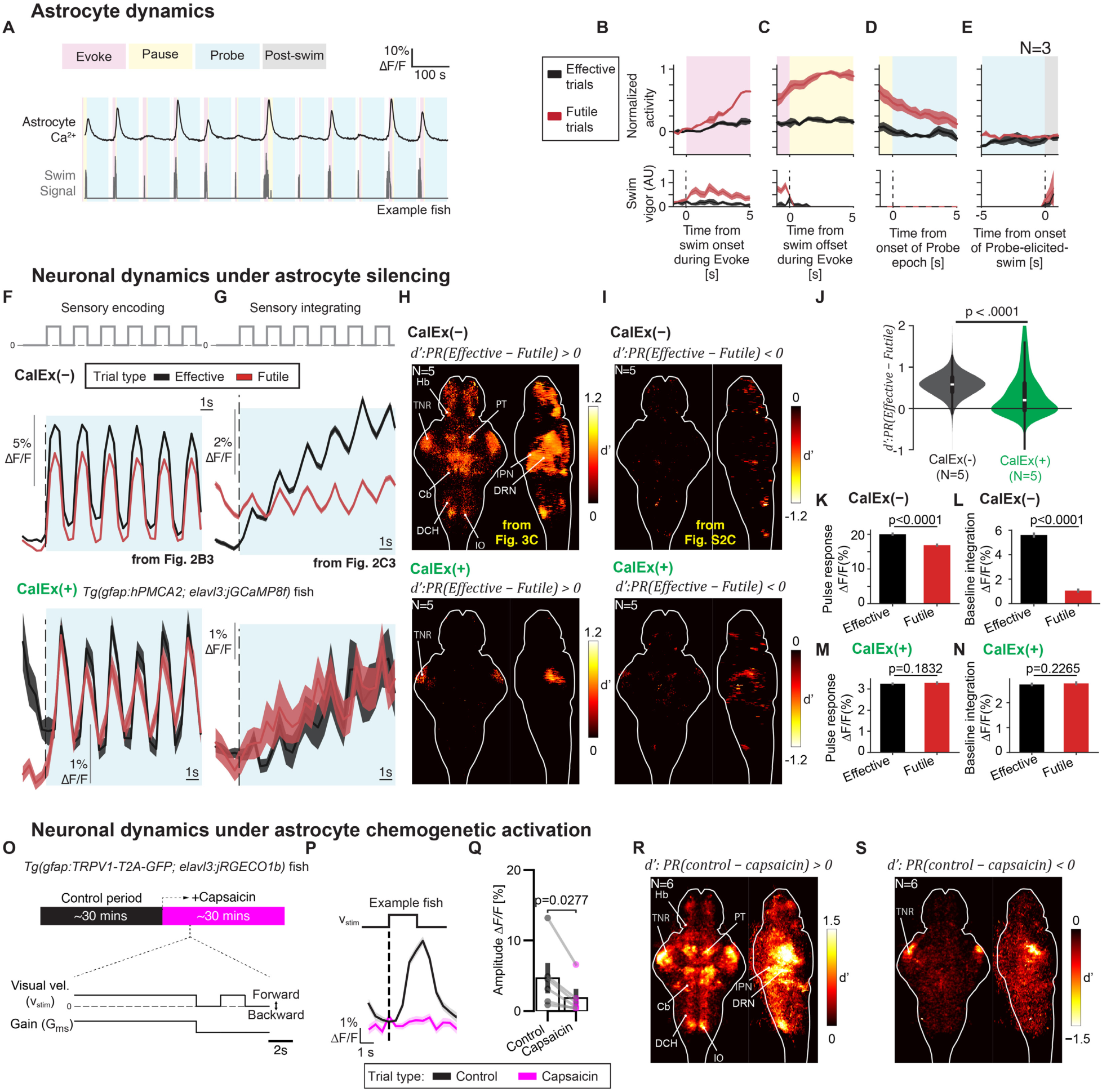
Astrocyte activation causally suppresses visual and integrative responses across the brain. **(A)** Example astrocyte activity during sequential behavioral tasks in one fish. Average brainwide astrocyte Ca²⁺ activity (top) and swim vigor (bottom) are shown. **(B-E)** Average astrocyte activity in Futile and Effective trials. Astrocyte calcium increased due to futile swims and such activation was maintained until probe-evoked swims. Shaded area, standard deviation across N = 3 fish. **(F-I)** CalEx-based silencing of astrocytic calcium eliminates the visual suppression after futile swims in *Tg(gfap:hPMCA2; elavl3:jGCaMP8f)* fish (CalEx(+) fish), in which futile swims can no longer effectively activate astrocytic calcium dynamics. Compared to the normal CalEx(−) fish (top), CalEx(+) fish (bottom) showed indistinguishable sensory responses **(F)** and sensory integration **(H)** during the Probe epoch after closed- or open-loop Evoke. They also showed less biased and weaker d’ brainwide **(H-I)**. N = 5 normal fish and N=5 CalEx(+) fish. **(F)** Average dynamics of top 10% sensory encoding neurons; **(G)** average dynamics of top 10% sensory integrating neurons. **(H)** Positive d’ map; **(I)** negative d’ map. **(J)** Distributions of d’ in normal, CalEx(−), (black) and CalEx(+) fish (green). Normal fish showed suppression of visual responses but visual responses in CalEx(+) fish were on average not suppressed in Futile trials; p<0.0001; χ^2^ test of the two distributions. **(K-L)** Both single pulse responses **(K)** and integration of pulses **(L)** were suppressed in normal fish on Futile trials. p<0.0001 for both, paired sign rank test across neurons in N=5 normal fish. **(M-N)** Single pulse responses **(M)** and integration of pulses **(N)** were indistinguishable between Effective and Futile trials in CalEx(+) fish. p>0.05 for both, paired signed-rank test across neurons in N=5 CalEx(+) fish. **(O-S)** Chemogenetic activation of astrocytes dampens neural responses to visual motion. **(O)** Experimental paradigm in *Tg(gfap:TRPV1-T2A-GFP; elavl3:jRECO1b)* fish with astroglial calcium influx induced by capsaicin. Bottom, experimental paradigm of a single trial. **(P)** Average pulse response in example fish across neurons in sensory-responding regions. Black, control trials; magenta, capsaicin trials; shaded area, SEM across trials. **(Q)** Bar plot of pulse response amplitude, N=6 fish. Black, control trials; magenta, capsaicin trials; error bar, SEM across fish; circles, single fish; two-tailed Wilcoxon signed-rank test. **(R)** Positive d’ map of the area under curve ΔF/F in control versus capsaicin trials for the 4 second period from probe onset; **(S)** negative d’ map; N=6 fish. See also Figure S7 for neuromodulator dynamics.

### Astrocyte activation is required to suppress the visual response

Astrocyte calcium dynamics, initially activated by futile swimming, closely tracked the emergence and decay of visual suppression that was correlated with the Disengaged state (Fig. 5A-E).^27^ To determine if astrocytes were the causal drivers of the changes in brain dynamics seen in the Disengaged state, we used CalEx (also known as hPMCA2; a human plasma membrane calcium ATPase pump that constitutively extrudes calcium) to reduce astrocyte calcium elevations and assess whether these transients were necessary for the suppression of visual responses following futile swimming.^28,54,64^ We measured sensory and sensory-integrative dynamics during Effective and Futile trials in fish expressing jGCaMP8f in neurons and CalEx in astrocytes.^65^ Astrocytic calcium responses evoked by futile swimming are attenuated in these ‘Astro-CalEx’ fish.^37^ We found that in Astro-CalEx fish, the difference of visual responses in Effective and Futile trials were statistically indistinguishable, both for sensory encoding (Fig. 5F) and for sensory integration (Fig. 5G); overall, brainwide state-dependent modulation was close to zero in CalEx fish (Fig. 5H-N). This result suggests that astrocyte calcium responses are necessary for internal-state dependent modulation of sensory processing across the brain.

To determine whether activation of astrocytes is sufficient to drive the suppression of visual responses, we used chemogenetics to selectively increase astroglial calcium levels brainwide, through expression of the rat TRPV1 receptor under the *gfap* promoter, which causes calcium influx upon activation by capsaicin (note that the native fish TRPV1 is insensitive to capsaicin) (Fig. 5O).^66^ Application of capsaicin suppressed sensory responses to forward motion pulses in fish expressing jRGECO1b in neurons (Fig. 5P,Q), resembling the brainwide modulation of sensory processing that occurred during the naturally induced Disengaged state (Fig. 5R,S, compared to Fig. 5H,I).^67^

These loss- and gain-of-function experiments show that astrocytes provide crucial modulatory control, enabling transient reconfiguration of brainwide computation to meet changing demands.

## Discussion

Unraveling the mechanistic origins of internal states is essential for understanding phenomena ranging from behavioral flexibility to mood fluctuations and maladaptive psychological states. A comprehensive mechanistic understanding of the underlying circuit changes has until now been difficult to obtain because neural recordings typically sample only small regions or fractions of the brain and restricted cell types, whereas internal states may be distributed, global, and involve complex intercellular interactions. This mismatch in scale has limited studies of their downstream impact, though important insights have come from highly accessible nervous systems,^4,20,29,63,68^ distributed electrical recordings,^69^ and the study of multicellular interactions in vivo and in brain slices.^27,28,37,54,55,58,70–73^ Furthermore, the true effects of internal states are difficult to measure as they are often coupled with ongoing behavioral changes. Here, by using whole-brain imaging and cellular manipulation and an assay that dissociates internal state from ongoing behavior, we showed that astrocytes promote behavioral flexibility by reconfiguring network dynamics, especially the properties of information accumulation over time, and reshaping sensory-motor transformations. Although behavioral unresponsiveness during the Disengaged state could, in principle, have arisen from the modulation of any stage in the sensorimotor pathway, we found that the internal state impinges in a coordinated manner on all stages of the sensorimotor transformation. This modulation has differential effects depending on the precise computation that a neuron partakes in, with strongest impact observed on the temporal integration stage, synergistic with direct motor suppression.^27,37,74^ Temporal stimulus integration lies between sensation and action, unconstrained by sensory organs or motor systems, yet central to sensory-to-motor transformation. This suggests an organizing principle for the way certain internal states reorganize behavior, by most strongly impinging on intermediate levels of sensorimotor computations and adjusting how these integrate different sources of information over time, to reconfigure pathways from sensation to action.^75^

The sensory-to-motor transformation model used here to conceptualize the data was, by necessity, simplified, and understanding the full pathway between sensory input to motor output requires more investigation, in particular to resolve the synaptic connections between these disparate brain regions.^53,76,77^ The cellular mechanisms by which neural circuit activity changes, such as the induction of increased leakiness in networks capable of integration and persistent activity,^78^ remains an open question. Our astrocyte manipulation experiments, including calcium extrusion using CalEx, lack spatiotemporal specificity, motivating the development of more physiological and spatially and temporally-precise astrocyte manipulation tools. Astrocytic release of ATP and extracellular conversion into adenosine has been shown to be a communication pathway from neurons to astrocytes in multiple species.^37,54,55^ Future work will determine whether this purinergic molecular pathway underlies the brainwide modulation discovered here in its entirety or if other mechanisms contribute in specific brain areas, and what molecular pathways within neurons mediate their effects on cellular processing.^79–81^

Futility-induced passivity may have evolved to preserve metabolic resources when actions fail to produce their desired outcomes and consume energy unnecessarily. A conceptual extension applies to neural firing, which is metabolically costly and produces damaging reactive oxygen species in neurons. Additionally, at the computational level, reducing firing may free resources for other computations or enhance signal-to-noise for encoding critical signals.^54^ Internal states are related to, but distinct from, behavioral states, which can be identified from observable behavior.^1^ We found that although behavior transitioned abruptly, brain dynamics evolved more smoothly and gradually between different internal states, highlighting that behavior is a partial readout of internal states and neurophysiological characterization is required to link the two.^82^ Beyond futility-induced passivity, internal states are diverse and shape behavior and physiology in distinct ways, and the approach used here—large-scale recording during a state-inducing paradigm and a multi-second integration and decision task—could be extended to such other states such as those driven by fear, hunger, or social context, and to multiple decision processes like choices to flee or freeze, explore or hide, or engage or avoid.^6,20^ Ultimately, studying a repertoire of states—including mixed and interacting states—within the same animal will clarify the full repertoire of the cellular, molecular, and network mechanisms by which past experience shapes future actions through persistent, coordinated modulation of brainwide computation.

## Acknowledgements

We thank Erik Snapp, the Johns Hopkins Neuroscience Training Program, and the Janelia Visiting Scientist Program for support. We thank Brett Mensh, Yu Mu, Florian Engert, Takashi Kawashima, Edmund Talley, Robert Johnson, Paul Tillberg, Liangyu Tao, Yuxin Tong, Peixiong Zhao, and Anoj Ilanges for comments on the manuscript. We thank members of the Ahrens lab, Jeremiah Cohen, Joshua Dudman, and Daniel O’Connor for their feedback on the project. We thank the Janelia Vivarium staff, Janelia Experimental Technology for infrastructural support for this project. This work was supported by Howard Hughes Medical Institute (M.B.A., J.E.F.), A*STAR National Science Scholarship (PhD) (J.-X.L.), the Simons Foundation Simons Collaboration on the Global Brain Award 542943SPI (M.B.A.), the National Institute for Theory and Mathematics in Biology through the National Science Foundation (grant number DMS-2235451) (J.E.F.), the Simons Foundation (grant number MPTMPS-00005320) (J.E.F.), National Science Foundation Award No. ECCS2032649 (B.B.), and National Institutes of Health (NIBIB) EB037653 (D.E.B.). This manuscript is the result of funding in part by the National Institutes of Health (NIH, EB037653) and subject to the NIH Public Access Policy. The NIH has been given a right to make this manuscript publicly available in PubMed.

## Author contributions

Conceptualization: J.-X.L., Z.W., D.E.B., J.E.F., M.B.A.; methodology: J.-X.L., Z.W., J.E.F., M.B.A., S.N., Y.Z., J.H., I.K., J.Z., L.L.L.; investigation: J.-X.L., Z.W., M.B., X.Y.; visualization: Z.W., J.-X.L.; data processing and analysis: Z.W., J.-X.L., A.S., B.B., X.M., W.Z., M.R., G.Y.; computational modeling: Z.W., M.B.A.; funding acquisition: J.-X.L., B.B., J.E.F., M.B.A.; project administration: M.B.A.; supervision: C.W., C.Z., D.E.B., J.E.F., M.B.A.; writing – original draft: J.-X.L., Z.W., M.B.A.; writing – review & editing: J.-X.L., Z.W., D.E.B., J.E.F., M.B.A..

## Supplemental Figures

**Figure S1.**
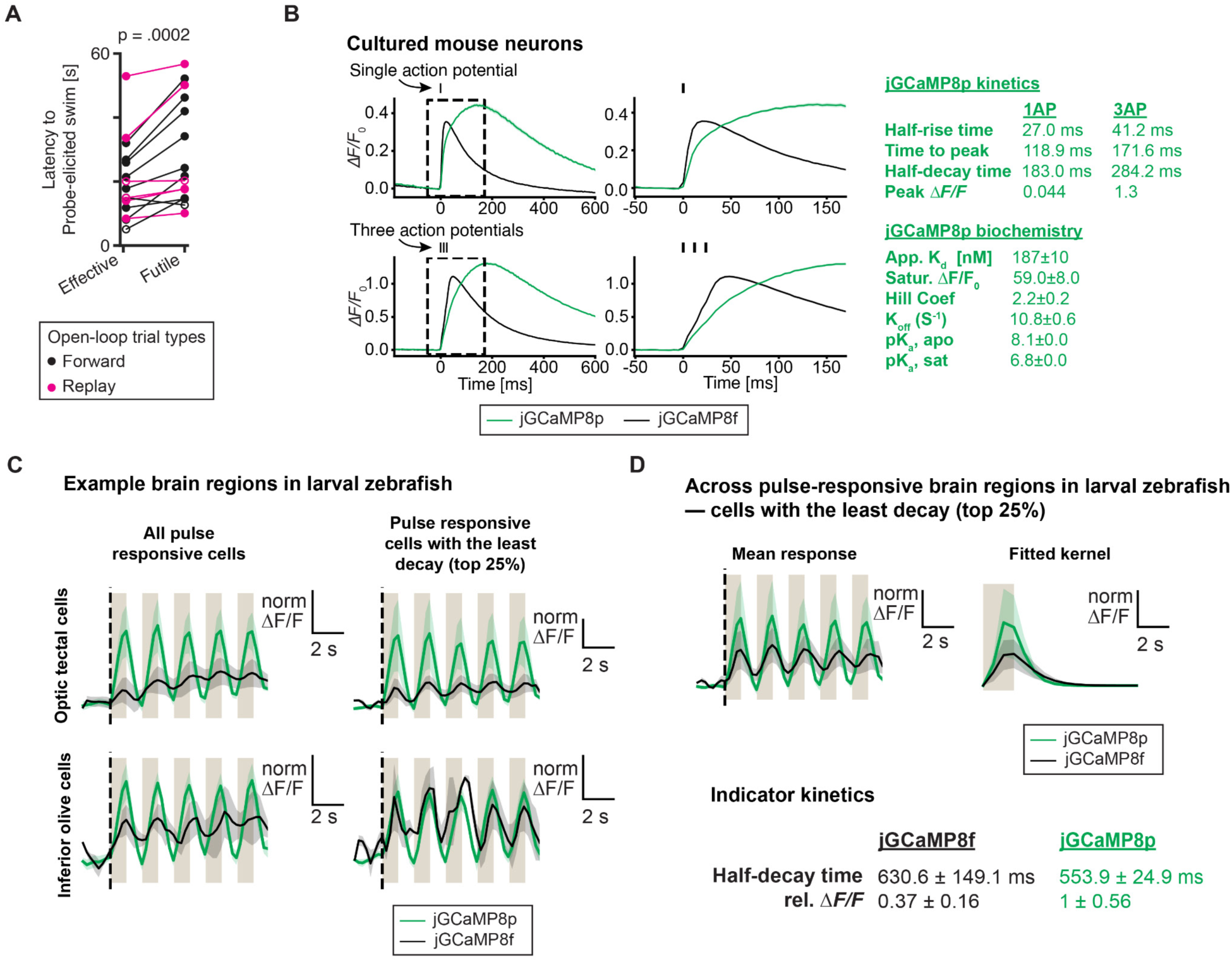
Related to Fig. 1. Statistics of fish behaviors in Effective and Futile trials, and calcium indicator kinetics. **(A)** Support material for Fig. 1G, with Futile trials broken down into two types – black: fish experienced *G_ms_=0* and *v_stim_* is a constant in Futile trials; magenta: fish experienced *G_ms_=0* and *v_stim_* is a replay from a previous Effective trial, i.e. open-loop replay, in Futile trials. Probe-evoked swims were delayed in Futile trials, ***p=0.0002, N=15 fish; p=0.0078, N=8 fish in constant-forward-grating Futile trials; p=0.0156, N=7 fish in open-loop-replay Futile trials; paired signed-rank tests. **(B)** Fluorescence responses of jGCaMP8p-expressing cultured mouse neurons to one (top) and three (bottom) action potentials in response to electric field stimulation in comparison to jGCaMP8f.^65^ Middle column, zoomed-in from the left (dash-boxed) to highlight rise kinetics (middle). Solid lines, mean; shaded areas, SEM; jGCaMP8p, green N=940 neurons, 7 plates, 40 wells; jGCaMP8f, black, N=804 neurons, 5 plates, 54 wells). Right, a summary table of jGCaMP8p kinetics. **(C)** *In vivo* fluorescence responses of all (left) and the fastest (right) neurons in the optic tecum (top) and inferior olive (bottom) expressing either jGCaMP8f (black) or jGCaMP8p (green) while fish is in the Disengaged state, aligned to the start of the Probe epoch (dotted line). Solid lines, mean; shaded areas, SEM across fish (jGCaMP8p, green, N = 4 fish; jGCaMP8f, black, N = 3 fish). Khaki bars, visual motion pulses. Vertical scale bar, normalized ΔF/F, representing that of the indicator with the larger ΔF/F, jGCaMP8p, for comparison. **(D)** *In vivo* fluorescence responses of sensory neurons with the least amount of motion integration, to isolate GCaMP8f and GCaMP8p kinetics for neuronal integration. Fish expressed either jGCaMP8f (black) or jGCaMP8p (green) and were in Futile trials. Left, responses to a series of visual motion pulses aligned to the onset of the Probe epoch (dotted lines). Right, the fitted impulse response to a single motion pulse. Khaki bars, visual motion pulses. Bottom, characterization of the kinetics of the impulse response. ΔF/F is normalized by that of the indicator with the larger ΔF/F, jGCaMP8p, for comparison. Note that in larval zebrafish at room temperature, jGCaMP8p exhibits faster kinetics than jGCaMP8f, whereas in mouse neuron culture jGCaMP8f exhibits faster kinetics, suggesting species/temperature differences. Due to GCaMP8p’s faster kinetics in zebrafish, this is the indicator used for most experiments in this work.

**Figure S2.**
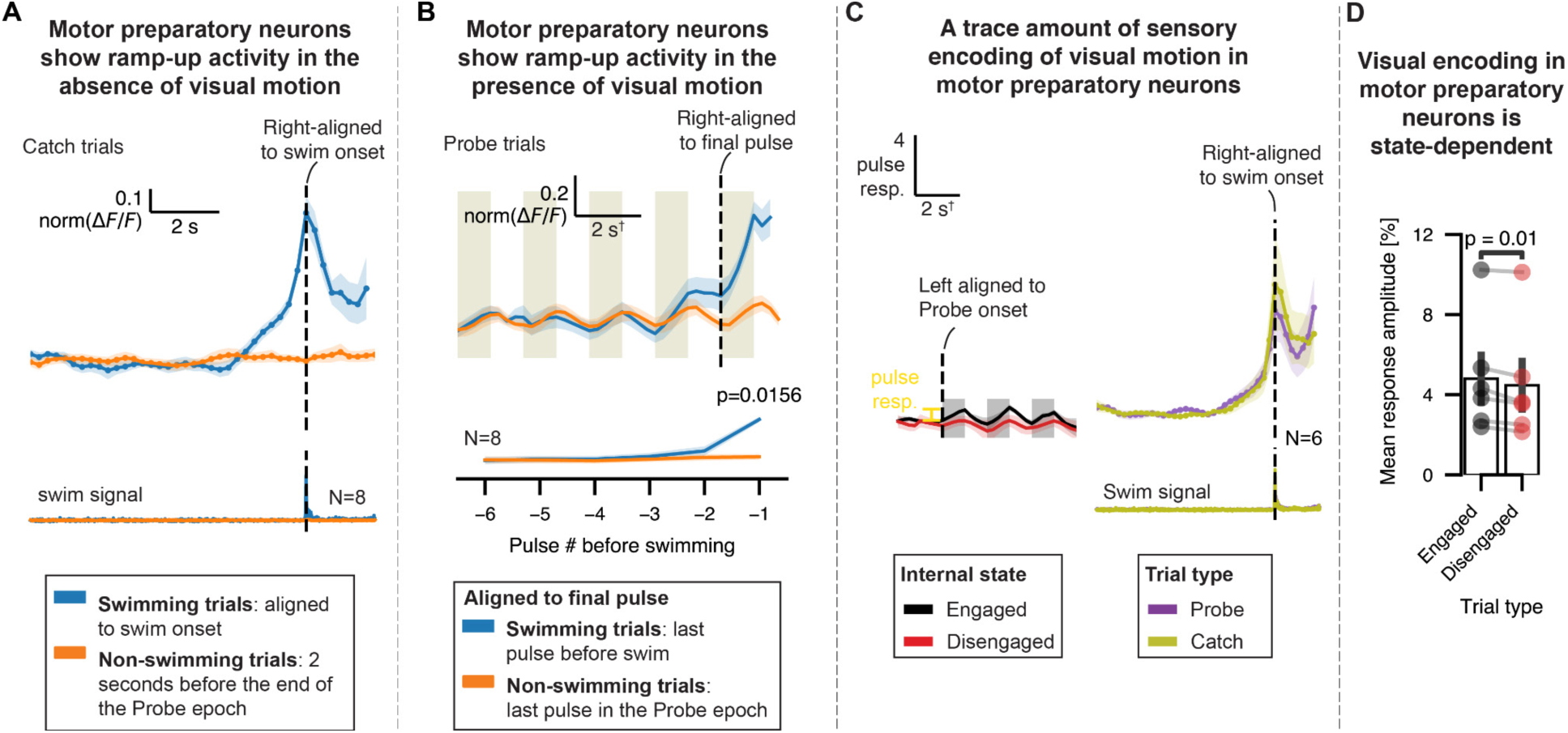
Related to Fig. 2E. Dynamics of motor preparatory neurons. **(A)** Motor preparatory neurons ramp up prior to swimming in Catch trials. Top, average neuronal activity ramps up from ∼3 s before swimming (blue; swimming trials); this ramping dynamics was not present in non-swimming trials (orange) (mean±sem across N=8 fish; min-max normalized to the lowest and highest activity values of the activity trace on swimming trials). Bottom, example swim signal from N=8 fish. **(B)** Motor preparatory neurons also exhibit weak visual motion response in Probe trials. Motor preparatory neurons displayed increased activity right before swimming. Top, the same convention as (A), except aligned to the last pulse before swimming, with traces min-max normalized to average activity trace of swimming trials. Khaki bars, visual motion pulses. Bottom, average neuronal response to visual pulses; line and shaded area, mean ± sem across N=8 fish; min-max normalized to the lowest and highest response; two-tailed Wilcoxon signed-rank test. **(C)** Motor preparatory neurons exhibit state-dependent visual response and ramp prior to swimming. Left, early-phase motor preparatory neuronal activity aligned to the onset of Probe epoch. Black, Engaged state; red, Disengaged state. Right, motor preparatory neuronal activity aligned to swimming; purple, Probe trials; olive, Catch trials. Shaded area, SEM across N=6 fish (two fish with insufficient state-related trials were excluded). **(D)** Early-phase visual response in motor preparatory neurons is state-dependent. Bar plots of the neural activity in the first 3 pulses of the Probe epoch in Engaged and Disengaged states; error bars, SEM across N=6 fish; two-tailed Wilcoxon signed-rank test. Across panels, s^†^ means 1 second for all but one fish; for one fish the visual motion pulses were 2 s long, interspersed by 2 s, in which case the time traces were scaled so that s^†^ represents 2 seconds to enable population averaging.

**Figure S3.**
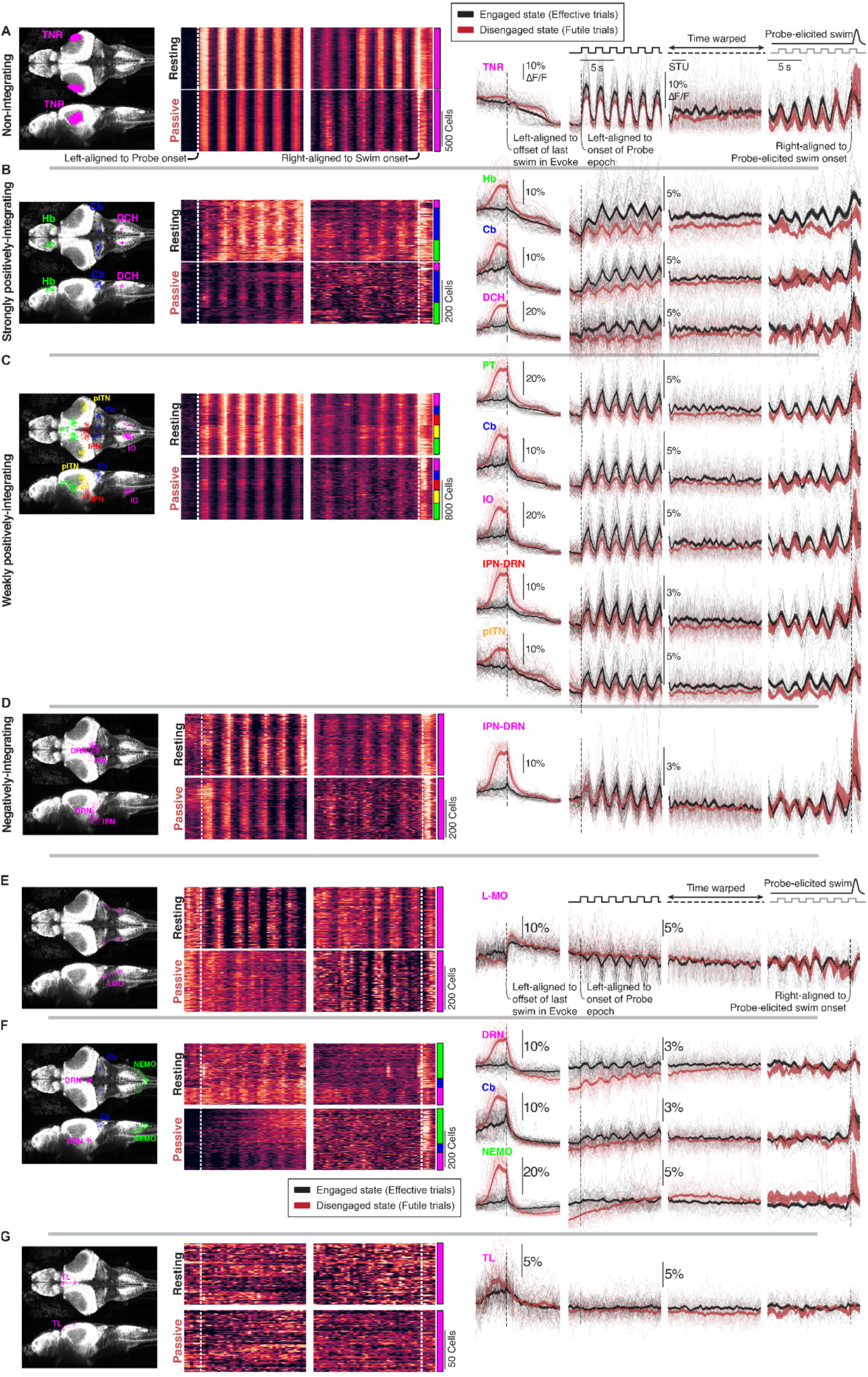
Related to Fig. 3. Multiple sensory regions are modulated during the transition into and by the Disengaged state in different ways. Details of the spatiotemporal clustering algorithm: state-modulated visual motion responsive cells (in a representative fish) are subdivided into groups of regions based on their integrative properties using hierarchical clustering of neural responses to the pulse series (a concatenated vector of time series of the neural activity in first 6 pulses in Effective and Futile trials;rows) and then by anatomical regions (DBSCAN subclustering^53^ based on neuronal spatial locations, with maximum distance between two neurons in the cluster <30 μm; sub-rows). The location of the anatomical regions in each group can be found in the brain maps (first column). Their single-cell activity can be found in heatmaps aligned to the onset of the Probe epoch (second column) and Probe-evoked swim (third column). Single-cell activity for both the Engaged and Disengaged states are shown. Single region activity is represented as traces aligned to the offset of swimming in the Evoke epoch (fourth column), aligned to the start of the Probe epoch (fifth column), aligned to both the start of the Probe epoch and onset of the probe-evoked swim (STU: scaled time unit) (sixth column), and aligned to the onset of the probe-evoked swim (seventh column). Single region activity in both the Engaged (black traces) and Disengaged (red traces) states are shown. Thin lines, neural dynamics on single trials; thick line,trial average; shaded area, SEM across trial. **(A)** The tectal neuropil region (TNR) is a visual-motion responsive region that does not integrate over pulses. The tectal neuropil has activity that is generally higher in the Evoke and Probe epochs, but that activity eventually converges between trial types before the onset of the Probe epoch. Activity during the Probe epoch is suppressed when the fish is in the Disengaged state, a trend that continues until the fish is about to swim. **(B)** The habenula (Hb), cerebellum (Cb) and dorsocaudal hindbrain (DCH) regions contain cells that strongly integrate over the visual motion pulses. All three regions are more strongly activated during the Evoke epoch before the fish enters the Disengaged state. This difference converges during the Pause epoch (note that this is a subset of the dorsocaudal hindbrain region that was clustered together with the other cell groups; other cells in this region show more persistent activity; Fig. S6). Activity during the Probe epoch is suppressed when the fish is in the Disengaged state. Activity in the habenula does not seem to converge between trial types, even after the fish performs the probe-evoked swim. On the other hand, activity in Cb and DCH regions seem to converge earlier. **(C)** The pretectum (PT), cerebellum (Cb), inferior olive (IO), interpeduncular nucleus / dorsal raphe nucleus (IPN-DRN), posterior lateral tectal neuropil region (pITN) contain cells that weakly integrate over the visual motion pulses. All regions are more strongly activated during the Evoke epoch before the fish enters the Disengaged state. This difference converges between trial types during the Pause epoch. Activity during the Probe epoch is suppressed when fish is in the Disengaged state and become increasingly less so during the course of the Probe epoch. Activity seems to have converged (i.e., became similar between trial types, but typically at a later time in Futile trials) before the onset of the probe-evoked swim. **(D)** The interpeduncular nucleus / dorsal raphe nucleus (IPN-DRN) region also contains cells that integrate negatively over the visual motion pulses. These cells are activated more strongly during the Evoke epoch before the fish enters the Disengaged state, a difference that converges during the Pause epoch. The visual motion pulses are encoded very similarly between the Engaged and Disengaged states. **(E)** The lateral medulla oblongata (L-MO) contains cells that encode the visual motion pulses negatively. Activity during the Evoke epoch is not very different between the Engaged and Disengaged states. However, the encoding of visual motion is suppressed to a large degree when the fish is in the Disengaged state. This response eventually recovers, but not until it gets closer to the Probe-evoked swim (note, this happens on average at later times in Futile trials). **(F)** The dorsal raphe nucleus (DRN), cerebellum (Cb) and NE-MO contains neurons that are much more highly activated during the Evoke epoch before the fish enters the Disengaged state. While that difference converges during the Pause epoch, presumably after the fish enters the Disengaged state, it eventually dips lower than that of the Engaged state. Much of this visual motion - independent difference is maintained until right before the fish performs the probe-evoked swim. **(G)** The torus longitudinalis (TL) is a region that does not respond to the visual motion stimuli but shows a difference in activity between the Engaged and Disengaged states. Specifically, activity is slightly suppressed when fish is in the Disengaged state, most clearly seen in the time warped traces.

**Figure S4.**
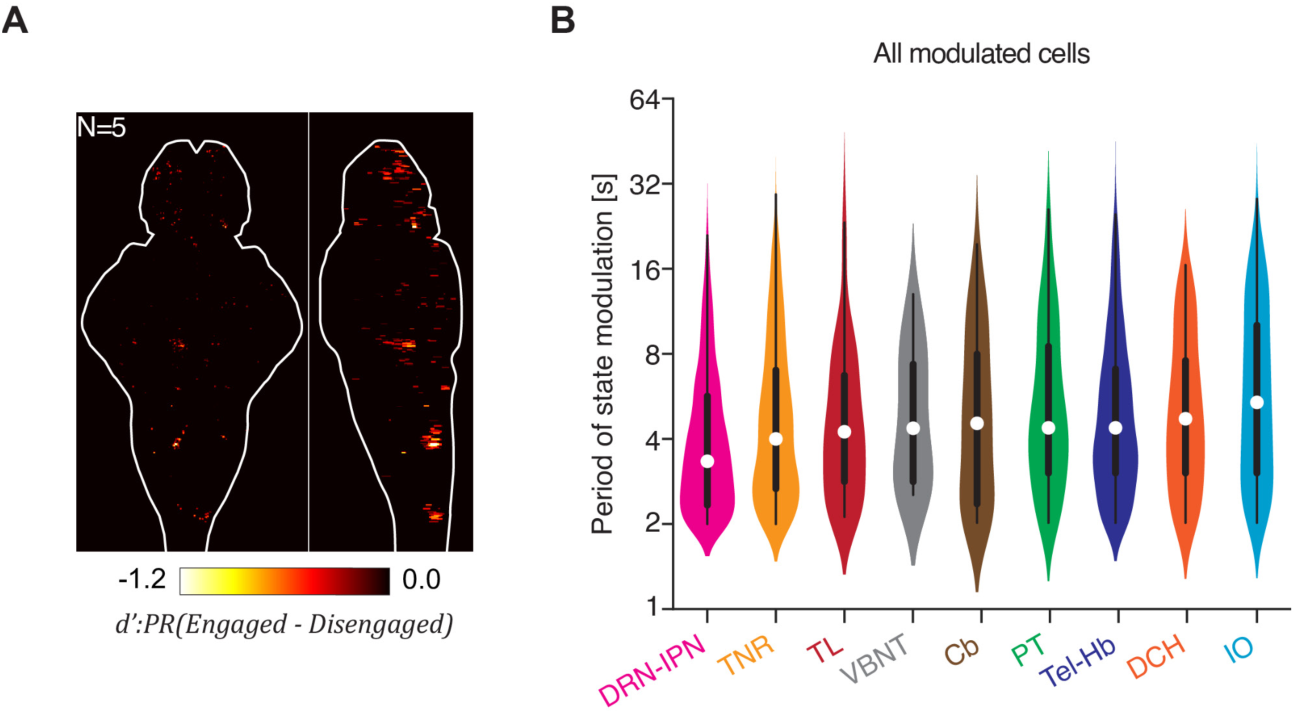
Related to Fig. 3. Additional analyses. **(A)** Support material for Fig. 3C. Brain map showing cells with a greater response in the Disengaged state than in the Engaged state, as quantified by the discriminability index (d’), with negative values here for comparison to positive values shown in Fig. 3C. Only 3.19 ± 2.01% of the pulse-responding neurons were negatively modulated; N=5 fish, mean ± SD. **(B)** Support material for Fig. 3G. Brain regions contain cells that are modulated for a large range of durations, from 0.7 sec to 42 sec. Violin plots showing the distribution of modulation duration across cells and across fish. A miniature box-and-whisker plot showing the quartiles of the data is also shown.

**Figure S5.**
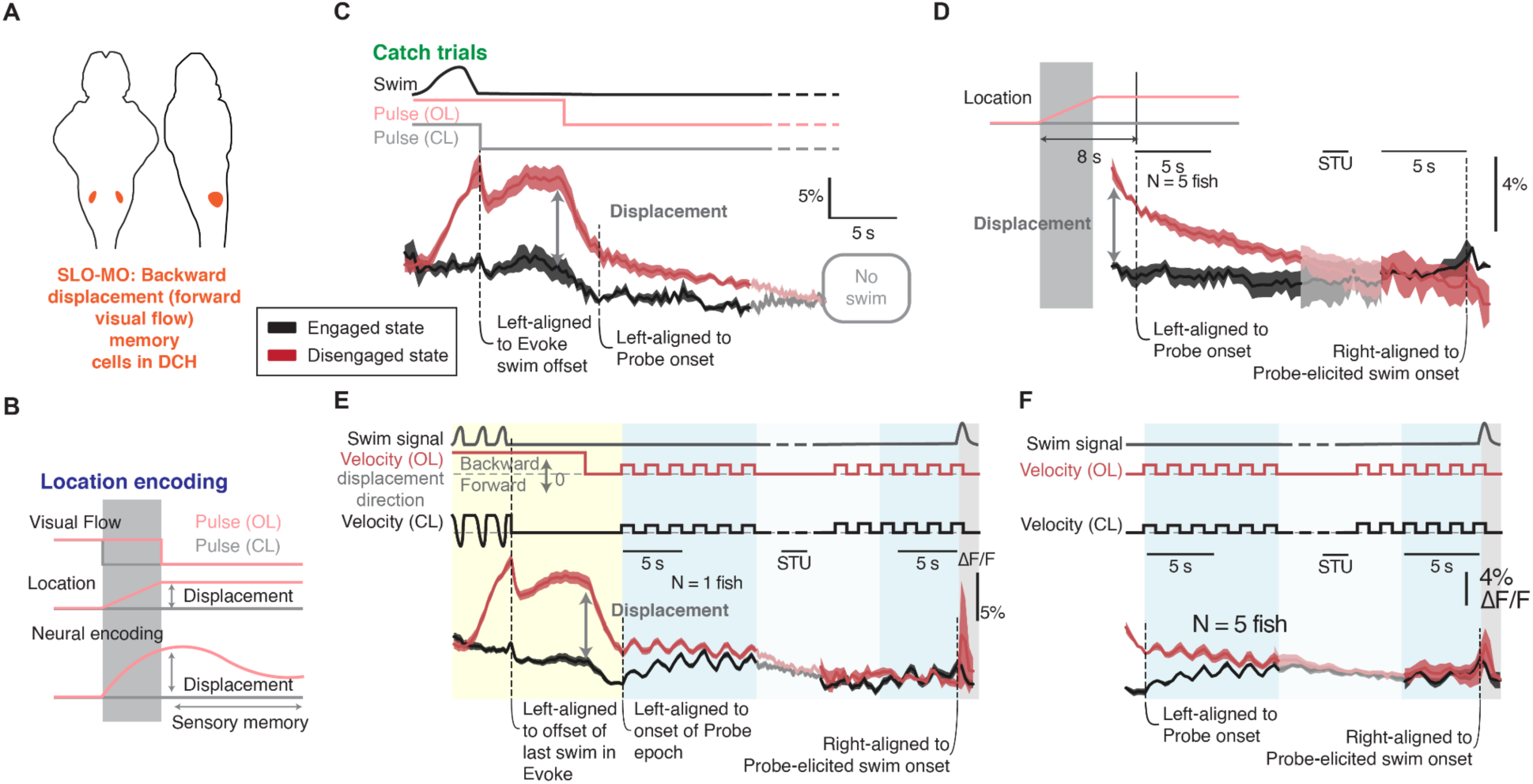
Related to Fig. 3. Location-encoding in SLO-MO dynamics. Due to the experimental design, backward displacement-encoding SLO-MO neurons showed higher activity in Futile trials, reflecting the additional ≥5 seconds of forward-grating exposure during the Evoke epoch. This finding is consistent with their proposed role in sensory integration and displacement memory.^41^ **(A)** Brain loci of the backward displacement-encoding cells in SLO-MO. **(B)** Schematics of visual flows and displacements during Effective (gray) and Futile (pink) types, where fish were exposed to forward-grating motion for additional 5 seconds during the Futile Evoke epoch, and the hypothesized neural encoding of displacement in SLO-MO. **(C-F)** SLO-MO activity is sensory history dependent and the difference of neural dynamics in Effective and Futile trials reflects the difference in displacements; black, Effective trials; red, Futile trials. **(C,D)** Average SLO-MO dynamics in Catch trials (no motion pulse during ‘Probe’ epoch). **(C)** Single fish; shaded area, SEM across trials. **(D)** Multiple fish; shaded area, SEM across fish, N=5. **(E,F)** the same as **(C,D)** in Probe trials. STU: scaled time unit.

**Figure S6.**
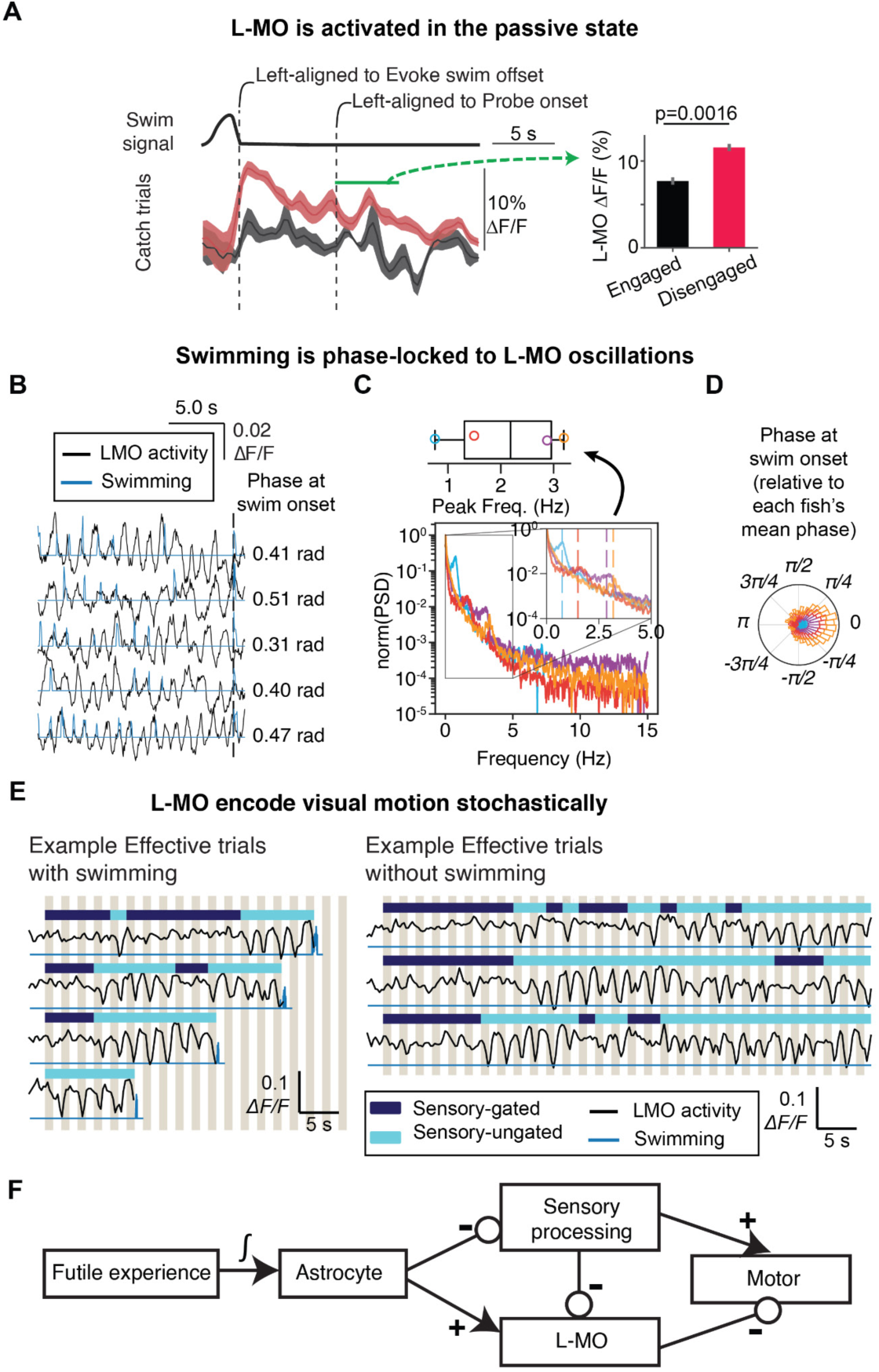
Related to Fig. 4. L-MO dynamics. **(A)** State-dependent L-MO dynamics in Catch trials. Left, neural dynamics aligned to the last swim in the Evoke epoch. Black, Effective trials; red, Futile trials; shaded area, SEM across fish, N=5. Right, Bar plots of neural dynamics in the early phase of Catch trials. Error bar, SD across neurons; p = 0.0016, paired signed-rank test across neurons. Previous study showed L-MO were excited during astroglial activation. We tested whether L-MO were excited in Futile catch trials. **(B)** Swimming is phase-locked to L-MO oscillations. Black, L-MO activity; blue, swim signal. Swimming phases to L-MO oscillations were shown for 5 example swim bouts on the right; vertical dotted line, swim onset. **(C)** L-MO oscillated at 0.5-3 Hz. Bottom, power spectral density of L-MO oscillations. Inset, zoomed in at <5 Hz. Top, box plot of peak frequencies of L-MO oscillations across N = 4 fish. Color, fish identity. **(D)** Rose plot of the histograms of swimming phase to L-MO oscillations across N=4 fish. Color, fish identity. **(E)** L-MO stochastically responds (navy, no response; light blue, response to visual motion) to visual motion (khaki bars) with decreased calcium in swimming trials (left) and non-swimming trials (right); response becomes robust before swim; black line, ΔF/F trace; blue line, swim trace. **(F)** Hypothesized model of astrocytic modulation of sensory processing and L-MO. Astrocytic calcium integrates futile swims. Previous work showed that astrocytes activate GABAergic neurons in area L-MO, a motor-suppression hub, thereby inhibiting swimming during the passive state.^27,37^ Reduced L-MO activity may relieve this suppression and facilitate swimming (Fig. 4F). This study shows that astrocytes also dampen sensory processing and integration (Fig. 5). In addition, forward visual motion inhibits L-MO activity (Fig. 4G, Fig. S6E), producing state-dependent disinhibition of L-MO when astrocytes are activated. Thus, astrocytic modulation of L-MO synergizes with their suppression of visual processing to reconfigure sensorimotor transformations during state switches.

**Figure S7.**
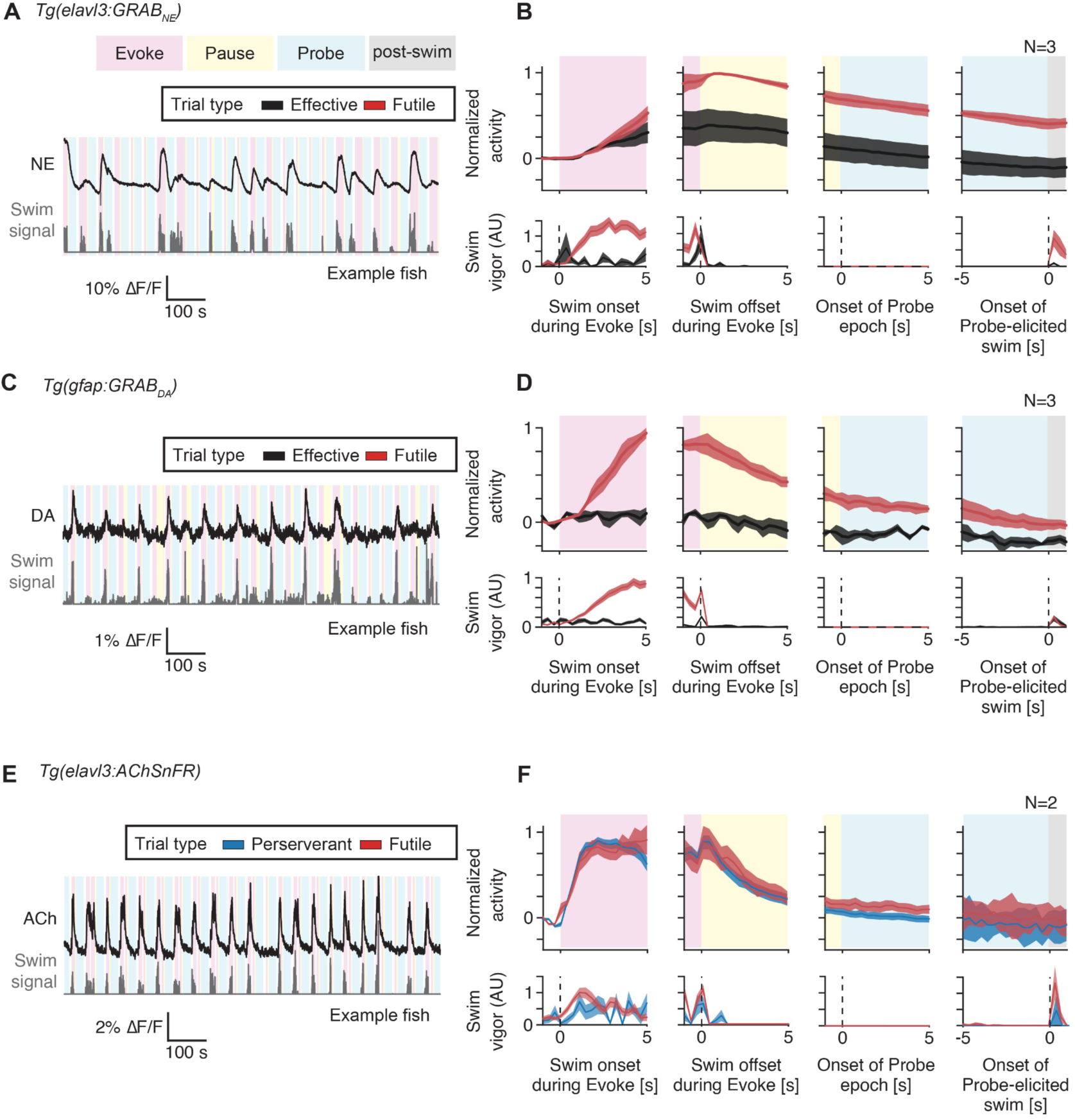
Related to Fig. 5. Futile swimming is associated with norepinephrine, dopamine and acetylcholine release. **(A,B)** Brainwide norepinephrine signals during trials. **(A)** Whole-brain averages show that norepinephrine levels rise sharply during strong futile swimming during the open-loop evoke in Futile trials. **(B)** Whole-brain averages show that norepinephrine is released to a large extent when the fish underwent strong futile swimming before entering the Disengaged state, with levels remaining elevated after the offset of swimming. Note that slow indicator dynamics at room temperature in zebrafish may contribute to the long time constants. Black, Effective trials; red, Futile trials. Shaded area, SEM across N=3 fish. **(C,D)** Brainwide dopamine release, conventions as in (A). Note that indicator dynamics at room temperature in zebrafish may contribute to observed slow dynamics. N=3 fish. **(E,F)** Acetylcholine release, depicted as in (A,B) but during Futile and Perseverant behavior, where fish underwent futile swimming but have yet to ‘give up’. The same comment about potentially slow indicator dynamics applies. N=2 fish.

## Methods

### KEY RESOURCES TABLE

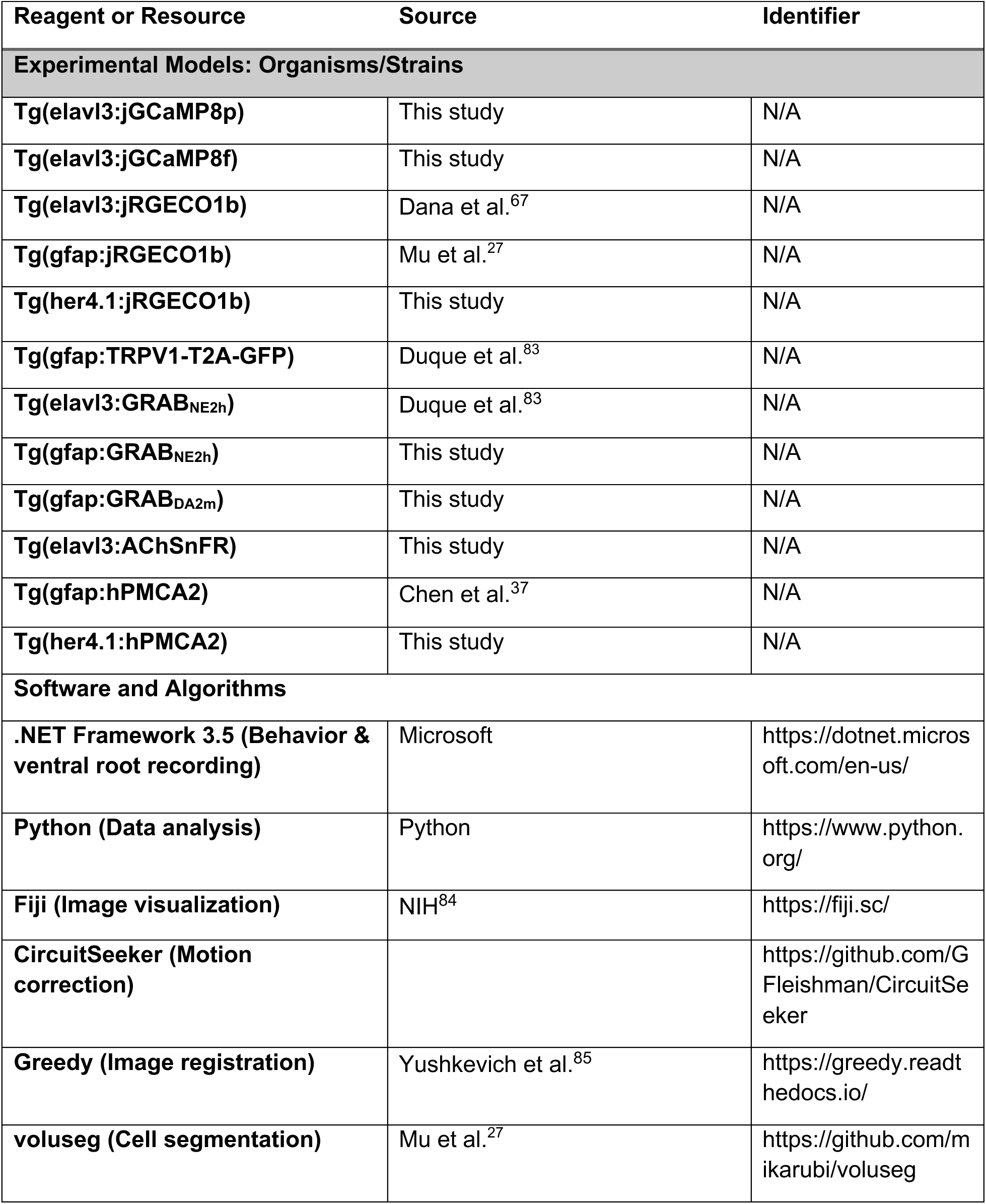
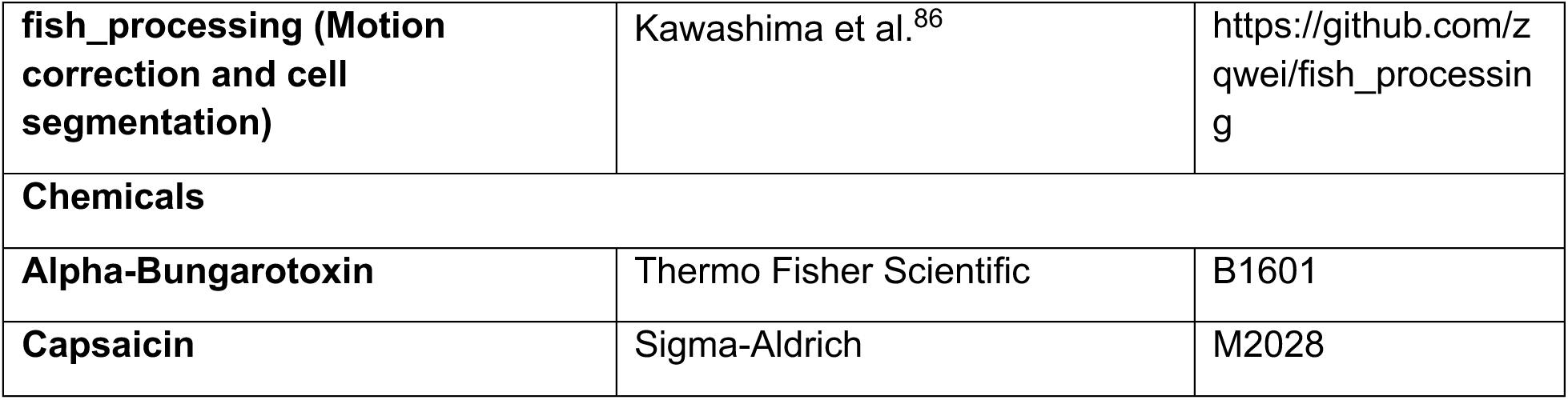

### EXPERIMENTAL MODEL AND STUDY PARTICIPANT DETAILS

#### Zebrafish husbandry

All experiments were conducted according to the animal research guidelines from NIH and were approved by the Institutional Animal Care and Use Committee and Institutional Biosafety Committee of Janelia Research Campus. Larval zebrafish were reared in Petri dishes at 0-3 days post fertilization (dpf) and 500 ml beakers thereafter at 28.5°C in 14:10 light-dark cycles.^87^ They were fed rotifers ad libitum from 4 dpf onwards. All experiments were conducted on 5-8 dpf larval zebrafish. Their sex could not be determined at this young age.^88^ All experiments were performed during daylight hours.

#### Genetically-encoded calcium sensor design

jGCaMP8p was developed during jGCaMP8 screening process, as described in Zhang et al.^65^ Briefly, the calmodulin (CaM) binding peptide RS20 in GCaMP6s was replaced with a peptide from endothelial nitric oxide synthase. The resulting variants, generated through structural-guided mutagenesis, were screened either in Escherichia coli or in cultured neurons. Among these, jGCaMP8p and jGCaMP8f were selected for their improved kinetics and sensitivity, and subsequently cloned into fish expression vectors. jGCaMP8p sequence:

MHHHHHHTRRKKTFKEVATAVKIIARLMGLKINVYIKADKQKNGIKANFHIRHNIEDGGVQLAYHYQQNTPIGDGPVLLPDNHYLSVESKLSKDPNEKRDHMVLLEFVTAAGITLGMDELYKGGTGGSMVSKGEELFTGVVPILVELDGDVNGHKFSVSGEGEGDATYGKLTLKFICTTGKLPVPWPTLVTTLTYGVQCFSRYPDHMKQHDFFKSAMPEGYIQERTIFFKDDGNYKTRAEVKFEGDTLVNRIELKGIDFKEDGNILGHKLEYNLPDQLTEEQIAEFKEAFSLFDKDGDGTITTKELGTVMRSLGQNPTEAELQDMINEVDADGDGTIDFPEFLTMMARKMKYRDTEEEIREAFGVFDKDGNGYISAAELRHVMTNLGEKLTDEEVDEMIREADIDGDGQVNYEEFVQMSTAK.

#### Transgenesis

Transgenic larval zebrafish were of either *casper* or *nacre* background. The transgenic line *Tg(gfap:jRGECO1b)* was generated in Mu et al.^27^ The transgenic line *Tg(elavl3:jRGECO1b)* was generated in Dana et al.^67^ The transgenic lines *Tg(elavl3:GRABNE2h)* and *Tg(gfap:TRPV1-T2A-GFP)* were generateed in Duque et al.^83^ The transgenic line *Tg(gfap:hPMCA2)* was generated in Chen et al.^37^ We generated for this study *Tg(elavl3:jGCaMP8p)*, *Tg(elavl3:jGCaMP8f)*, *Tg(her4.1:jRGECO1b)*, *Tg(elavl3:GRABNE2h)*, *Tg(gfap:GRABNE2h)*, *Tg(gfap:GRABDA2m)* and *Tg(elavl3:AChSnFR)* using the Tol2 transposon system.^89^

#### Transgenic zebrafish

For the imaging of neuronal calcium, we used the following transgenic zebrafish lines:

- *Tg(elavl3:jGCaMP8p)* (this paper)
- *Tg(elavl3:jGCaMP8f)* (this paper)
- *Tg(elavl3:jRGECO1b)*^67^

For the imaging of astrocytic calcium, we used the following transgenic zebrafish lines:

- *Tg(gfap:jRGECO1b)*^27^
- *Tg(her4.1:jRGECO1b)* (this paper)

For the imaging of extracellular norepinephrine, we used the following transgenic zebrafish lines:

- *Tg(elavl3:GRABNE2h)*^83^
- *Tg(gfap:GRABNE2h)* (this paper)

For the imaging of extracellular dopamine, we used the following transgenic zebrafish lines:

- *Tg(gfap:GRABDA2m)* (this paper)

For imaging of extracellular acetylcholine, we used the following transgenic zebrafish lines:

- *Tg(elavl3:AChSnFR)* (this paper)

For silencing astrocytes, we used the following transgenic zebrafish lines:

- *Tg(gfap:hPMCA2)*^37^
- *Tg(her4.1:hPMCA2)* (this paper)

For the chemogenetic activation of astrocytes, we used the following transgenic zebrafish lines:

- *Tg(gfap:TRPV1-T2A-GFP)*^83^

### METHOD DETAILS

#### Fish selection and sample preparation

Before imaging and fictive behavioral recording experiments were performed, larval zebrafish at 5-8 dpf were transferred from their home beaker into a wide petri dish (LabStock 150 mm x 30 mm clear borosilicate glass) where they were fed rotifers for at least 1 hour.

To prevent motion artifacts in the brain images we acquired, fish were placed in a droplet of the nicotinic acetylcholine receptor antagonist ɑ-bungarotoxin (1 mg/ml) for 5-10 seconds. ɑ-bungarotoxin solution was prepared by dissolving ɑ-bungarotoxin conjugates (Invitrogen) in external solution (134 mM NaCl, 2.9 mM KCl, 2.1 mM CaCl2, 10 mM HEPES, 10 mM glucose; pH 7.8; 290 mOsm). Following that, they were transferred to a dish with fresh fish water, where their motility and health were closely monitored. After fish stop swimming and exhibit reduced motility to tactile stimuli, normally after 10-15 minutes, they were transferred to a centered acrylic pedestal on a custom-fabricated sample holder^36^ (CAD design available upon request), where they were embedded in low-melting point agarose (Sigma-Aldrich Inc.). Lastly, the chamber was filled with fresh fish water.

#### Fictive behavior in virtual reality

We placed the fish in a virtual reality system^36^ to study their behaviors and brain dynamics in a newly designed sensorimotor task (Fig. 1D,E). In brief, 1-dimensional gratings (width 2.5 mm), moving along the axis of the fish, were projected onto a diffuser at the bottom of the chamber from the outside, ∼1 cm below the fish, using a miniature projector (Sony Pico Mobile Projector; MP-CL1). The velocity of the gratings, *v_stim_*, was defined by:

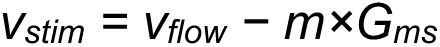

where *v_flow_* is an experimenter-defined forward-moving grating velocity; *G_ms_* refers to motosensory gain and is an experimenter-defined parameter modulating the change of velocity of the gratings per unit of motor drive; and *m* refers to motor drive and is the power computed from the firing of motor neuron axons in the ventral root.^90–92^

Agarose was removed around the tail to place electrodes on both sides of the tail for recording fictive swim signals originating from ventral root motor neuron axons.^90,93^ Electrodes were made with borosilicate pipettes (TW150-3, World Precision Instruments) pulled by a vertical puller (PC-10, Narishige) and shaped by a microforge (MF-900, Narishige) to have a final inner tip diameter of approximately ∼40 μm. Light negative pressure was applied to the electrodes to ensure a good seal between the glass pipette and the tail of the fish. Electric signals were amplified by a MultiClamp 700B microelectrode amplifier in current clamp and acquired using National Instruments DAQ boards. Signals were sampled at 6 kHz, and hardware band-pass filtered with a 3 kHz/100 Hz low-pass/high-pass cutoff.

Electrophysiological signals were acquired, and visual stimuli were presented using custom software written in C# (Microsoft).^90^ Swim vigor (*m*) was immediately decoded from this signal by computing its standard deviation within a sliding 10 ms window, and swim bouts were detected by setting an automated threshold, as described in ref. (*104*). Briefly, with *y* being the electrophysiological signal, with *x=√((y−y*ker_1_)*^2^*\*ker_2_)* where *ker_1_* and *ker_2_* are smoothing kernels on timescales of 100 ms and 20 ms, respectively, and * indicates convolution, *d(x)* the distribution of *x* values, *x_max_* was defined such that *d(x_max_)=max(d)* (the mode), *x_min_* was set to the highest value of *x* at which *d(x_min_)>0.01×d(x_max_)*, then the threshold was set to *th=x_max_+c(x_max_-x_min_)* with *c* set between 1.8 (lower noise recordings) and 5 (higher noise recordings).

To motivate the equation above (*v_stim_=v_flow_−m×G_ms_*), when fish move through their environment, the visual scene beneath them moves opposite to their fictive bodily motion, with a velocity we call *v_stim_*. For example, if fish swim forward, the visual scene beneath them accelerates backward in their field of view (*v_stim_* will become smaller or negative). When they are moved backward by a water current, the visual scene moves forward (*v_stim_>0*, tail-to-head motion) in their field of view. We used this relationship to simulate the effects of flow and swimming as fish were head-restrained in VR. When the whole projected visual scene (here represented by gratings) is moving forward relative to the fish (*v_stim_>0*, in the tail-to-head direction), the fish will have the sensation of being displaced backwards. This causes the fish to swim towards the direction of motion (i.e. forward) in order to maintain its position in space (i.e. positional homeostasis). To simulate forward movement during swimming, we bring the gratings towards the fish by moving the gratings backwards (i.e. *v_stim_<0*, in the head-to-tail direction).

The extent to which the gratings move away from the fish while it is swimming is tuned by *G_ms_* (Fig. 1B,C). When *G_ms_>0*, the gratings effectively move backwards, giving the fish the sensation of moving forward and towards its original location. This configuration is referred to as *closed-loop*, because the virtual environment moves in accordance to fish behavior. When *G_ms_=0*, grating velocity cannot be controlled by the fish, and the fish continues to be pushed farther away from its original location, regardless of its swim vigor. This is also referred to as *open-loop*.

After the ventral root recording “matures” (i.e. the swim signal is large enough to clearly discern swim bouts), which typically occurs in about 10 minutes, we tuned two parameters with the aim of reproducing freely-swimming optomotor response behavior. We tuned *v_flow_* so that forward visual motion (i.e. in the tail-to-head direction) reliably triggers the optomotor response. We also tuned *G_ms_* such that the gratings moved backwards (i.e. in the head-to-tail direction) whenever the fish swam in closed-loop. We visually confirmed that the fish could comfortably achieve positional homeostasis (i.e. remain in the same spot) over the time span of minutes. The fish fictively behaved in this virtual 1-dimensional track throughout an experiment.

Before every experiment, we verified that fish were stimulus responsive. Firstly, we checked that fish would swim in response to forward-grating visual motion and stopped swimming when switched to the static gratings. Secondly, we checked that fish would swim less vigorously at high *G_ms_* and more vigorously at low *G_ms_*, recapitulating known motor adaptation.^24,90^ Finally, we verified that fish behaved normally during brain imaging, since, in principle, the excitation laser could act as a source of distraction. By using minimal laser power, we ensured that fish still exhibited visual motion responsiveness and motor adaptation, and that the resulting swim patterns were similar in vigor and cadence to their non-imaging state.

#### Behavioral paradigm: Effective and Futile trials

On Effective trials, fish first swam in a closed-loop environment for 3-10 seconds (depending on basal swim rate of the fish) to ensure an Engaged state (*v_flow_>0*, *G_ms_>0*). This was followed by the Evoke epoch of another 5-15 seconds of closed-loop to maintain the Engaged state (*v_flow_>0*, *G_ms_>0*). This was followed by an 8 second Pause epoch (*v_flow_=0*, *G_ms_>0*) during which fish typically stopped swimming in a “quiescently Engaged” state. This was followed by the Probe epoch with 1 second pulses of forward visual motion alternating with 1 second periods of no motion, where *v_flow_* of the motion pulses was typically set to a lower value than *v_flow_* during the Evoke epoch (on average a factor of 2.5 slower) to ensure delayed responses. The Probe epoch was capped at 20-60 seconds. This was followed by a 2-10 second (depending on basal swim rate) inter-trial period (*v_stim_=0*, *G_ms_>0*), after which the next trial, i.e. a Futile trial, began.

On Futile trials, fish first swam in a closed-loop environment for 3-10 seconds to ensure an Engaged state (*v_flow_>0*, *G_ms_>0*). This was followed by the Evoke epoch in which fish swam in an open-loop environment (*v_flow_>0*, *G_ms_=0*) until fish entered futility-induced passivity, which was defined as 5 seconds of no swimming. Automatic online detection of futility-induced passivity, i.e. the Disengaged state, triggered a 3 second Pause epoch (*v_flow_=0*). This was followed by the Probe epoch as above, the inter-trial period as above, and the next trial, i.e. an Effective trial. The Pause epoch on Futile trials was set to 3 seconds to equalize the time of no-swimming before the Probe epoch across Effective and Futile trial types, which on both trial types was 8 seconds.

#### Light-sheet imaging

We removed agarose around the head for whole-brain imaging^94^ in a custom-built light-sheet microscope that enabled high-speed acquisition of images of the whole brain while fish fictively behaved in virtual reality, as described in Vladimirov et al.^36^ Briefly, images of the brain were acquired using two excitation lasers scanning the tissue at orthogonal angles – one entering the brain in between the eyes in the anterior-to-posterior direction (front laser) and the other entering the brain laterally (side laser). We used a laser with a wavelength of 488 nm for the excitation of jGCaMP8p and jGCaMP8f (calcium sensors), GRAB_NE2h_, GRAB_DA2m_, and AChSnFR (neuromodulatory sensors), and used a laser with a wavelength of 561 nm for the excitation of jRGECO1b (a calcium sensor used for astrocytes). Each of the illumination arms consisted of an air illumination objective (4x/0.28 NA, Olympus) mounted horizontally on a piezo stage, a tube lens, an f-theta lens and a pair of galvanometer scanners (Cambridge Technology). Collimated beams were scanned laterally and along the z-axis of the image space with the scan mirrors and f-theta lens. To minimize distractions from the visual stimulus beneath the fish and prevent the laser from directly stimulating the retina, the side laser was rapidly turned off at positions where the beam would enter the eye.

The detection arm consisted of a water-dipping objective (16x/0.8 NA, Nikon) mounted on a piezo stage (Physik Instrumente) vertically above the sample. The arm is followed by a tube lens and a sCMOS camera (Orca Flash 4.0, Hamamatsu). Fluorescence light from green fluorescent indicators (jGCaMP8p, jGCaMP8f, GRAB_NE2h_, GRAB_DA2m_, and AChSnFR) and red fluorescent indicators (jRGECO1b) were filtered by a band-pass filter (525/50 nm, Semrock) or a long-pass filter (590 nm, Semrock) respectively before they are acquired by the camera.

Stacks of either 300 µm or 400 µm were acquired per experiment at x, y, z voxel size of 0.406 um, 0.406 um and 5 µm, respectively, resulting in acquisition rates of 3 Hz or 2 Hz respectively. Experiments evaluating the effect of behavioral state on visual processing and behavior took ∼3-4 hours to run. Experiments evaluating the effect of chemogenetic stimulation of particular brain regions on visual processing and behavior took 1.5-2 hours to run. A high-resolution stack was acquired at x, y, z resolutions of 0.406 um, 0.406 um and 1 µm, respectively, either before or after the experiment to facilitate motion correction and registration of the data to brain atlases.

#### Silencing of radial astrocytes

To determine if astrocytes are necessary for the observed visual suppression, we characterized the effects of futility in fish expressing hPMCA2^64^—a human plasma membrane Ca^2+^ ATPase pump that constitutively extrudes Ca^2+^—in astrocytes. Similar to previous experiments, fish either experienced futile swimming in an open-loop environment or effective swimming in a closed-loop environment, before a series of forward-moving grating pulses were presented.

#### TRPV1-mediated activation of radial astrocytes

To determine if astrocyte activation is sufficient for the observed visual suppression, we presented fish with a single forward-moving grating pulse during and without the chemogenetic activation of astrocytes using *Tg(gfap:TRPV1-T2A-GFP)* fish whose astrocytes express the calcium-permeable channel TRPV1 which is activated by capsaicin. Capsaicin was prepared by dissolving capsaicin granules (Sigma-Aldrich) in ethanol (Sigma-Aldrich). Capsaicin was bath-applied via a 100-1000 µL mechanical pipette (Eppendorf) to a final concentration of 2 µM. The insides of the pipette were thoroughly rinsed by ejecting and injecting bath water ∼5 times, therefore also facilitating the mixing of capsaicin into the bath. The times where capsaicin was added was recorded. To allow for experimental conditions to equilibrate, that and a subsequent 2 minute period were excluded from further analysis.

### QUANTIFICATION AND STATISTICAL ANALYSIS

#### Trial classification

Trials were classified as follows:

- The Engaged state was defined (within Effective trials) as corresponding to trials where the fish swam continuously (with an inter-swim-interval lasting at most 3 s) until the end of the Evoke epoch in closed-loop (*G_ms_>0*). We excluded trials in which fish paused for >3 s at the end of the Evoke epoch.
- The Disengaged state was defined (within Futile trials) as corresponding to trials where fish swam at the beginning but stopped swimming for >5 s during the Evoke epoch, including at the end of the epoch.
- Perseverant trials were trials in which fish swam in open-loop but did not switch to a passive state and only stopped swimming upon the transition to the Pause epoch.

#### Pre-processing of whole-brain imaging data

All brain images were put through a pre-processing pipeline that consisted of three stages: motion correction, cell segmentation, and registration to a brain atlas.

Motion correction was performed with CircuitSeeker, a SimpleITK-based registration package developed for brain images (https://github.com/GFleishman/CircuitSeeker). SimpleITK^95^ is a programming interface to the algorithms and data structures of the Insight Toolkit,^96^ an open-source toolkit for multi-dimensional scientific image processing, segmentation and registration. CircuitSeeker has now been updated and merged into Bigstream (https://github.com/JaneliaSciComp/bigstream).

Cell segmentation was performed with voluseg (https://github.com/mikarubi/voluseg), a volumetric cell segmentation pipeline,^27^ and a second custom pipeline (https://github.com/zqwei/fish_processing) described in ref.^86^

Finally, images were registered to an anatomical brain (acquired using 1P confocal microscopic imaging) using a custom Python code based on Greedy (https://github.com/pyushkevich/greedy)^85^ to perform brain-map analyses across fish. To identify labels of brain regions, we registered the anatomical brain to Z-Brain Atlas (https://zebrafishexplorer.zib.de/).^97^

#### Analysis of state modulation along the sensory-to-motor pathway

For the analysis of the cells supporting the sensory-to-motor transformation of the optomotor response and their modulation by internal state, we selected fish that (1) were imaged using jGCaMP8p sensor at 3 Hz; (2) visual motion stimuli moved 1 second forward with a 1 second interval during the Probe epoch; (3) were complete and well-registered to the brain atlas; (4) whose neural response to visual motion has a good SNR (the percentage of visual motion responsive cells are around 16.92 ± 6.56% of the entire population, with a range of 11.19% - 27.19%). We selected the trials where fish swam during the Evoke epoch and did not swim during the Pause epoch. For Effective trials, we also required that the fish stopped swimming at most 3.3 s before the end of the closed-loop Evoke epoch, beyond which we consider the fish to have entered into a Disengaged state. In analysis with trials aligned both to the Probe onset and to the probe-elicited swims (Fig. 4A,B,G, Fig. S3, S5), we excluded the trials with <20-second latency to probe-elicited swims to reduce the overlap of the periods between the two alignments.

#### Visual motion-responsive cells

We computed the correlation between neuronal activity and the visual motion pulse series (*v_stim_*) during the early phase of the Probe epoch (the first 6 pulses, i.e. 12 seconds) using Spearman’s rank correlation in non-swimming trials (Fig. 2B,D). Visual-encoding cells were chosen as p<0.05 for correlation. Multiple-comparison correction was not performed because the final statistics were performed across fish instead of across neurons.

#### Integration time constants of visual motion-responsive cells

We characterized the integrative time constants, *τ*, of the visual response (Fig. 2C) for each visual motion-responsive cell, by fitting the neural dynamics ΔF/F(t) from visiual motion series *v_flow_(t)* as:

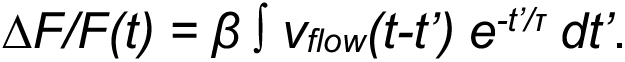

*β* and *τ* were fit by minimizing the squared difference between the model prediction and actual neural response, and (given that the indicator decay of GCaMP8p time constant is only ∼0.5 seconds and can therefore be ignored) *τ* is reported as the neuronal time constant.

#### Leakiness of integration of visual motion-responsive cells

We characterized the level of integration, i.e. *1/leakiness* or *leakiness^-^*^1^, of the visual response (Fig. 2) for each visual motion-responsive cell, as the ratio of ΔF/F after the sixth pulse to that from a perfect integrator based on the sum of the amplitudes of first six pulses:

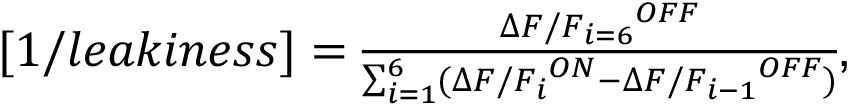

where ΔF/F^ON^ and ΔF/F^OFF^ is the ΔF/F at the last point during pulse on and off, respectively.

#### Detection of motor preparatory cells

Motor preparatory neurons were defined as those that ramp up ahead of probe-evoked swims on single trials (Fig. 2E). To search for activity ramps without biasing the analysis to any particular ramp duration, we computed Spearman’s rank correlation between the neural dynamics and the latency to swim in variable durations (5, 10, 15, and 20 s). When we found out that 5 s produced on average the highest *ρ*, we performed another local search at ramp times of 3, 4, 5, 6, and 7 s. Finally, based on the values of *ρ* computed using 3-7 s intervals, we defined motor preparatory cells as any cell had a significant positive correlation (top 5% of cells with *ρ*>0; p<0.05) for all of those durations.

#### Detection of motor-related cells

We computed the motor correlation of each neuron as Spearman’s correlation between the neural dynamics and swim vigor (*m>0*) during open-loop *evoke* (where *v_stim_=v_flow_* was constant). Motor cells are those with p<0.05 Spearman’s correlation. Correction for multiple comparisons was not performed because the eventual statistics were done not on neurons but on fish.

#### Two-way ANOVA in comparisons

We performed two-way ANOVA for comparison based on trial types (Effective or Futile) (Fig. 2E4, 2F4), where a measure can be modeled as:

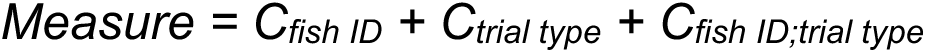

where *C_fish ID_* is a fish-identity-dependent constant; C_trial type_ is a trial-type-dependent constant; *C_fish ID;trial type_* is a constant that depends on both. We reported p values for *C_trial type_* in the main text. For both Fig. 2E4 and 2F4, p<0.0001 for *C_fish ID_*; p>0.1 for *C_fish ID;trial type_*.

#### Degree of modulation

We measured degree of modulation using the discriminability index,

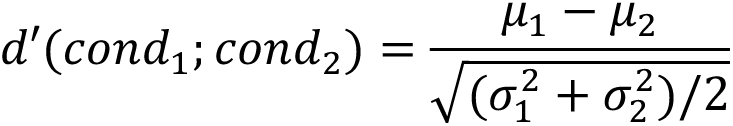

where *μ_1_* and *μ_2_* are the means of condition 1 and 2, respectively; *σ_1_* and *σ_2_* are the standard deviations of condition 1 and 2, respectively. In measure of the degree of modulation (Fig. 3), condition 1 and condition 2 are Effective and Futile trials respectively and we use the area under the curve of the neural dynamics of single pulses (starting from each pulse onset) in the first 6 pulses in this computation.

#### Timing of probe-evoked swim predicted by neural dynamics

We used the neural responses from the behavior-predictive sensory regions (Fig. 4C) to the n^th^ pulse before probe-evoked swim (0<n<8; t seconds before swimming) to predict its latency to the swim (t seconds). We fitted this neural data predictor using Gradient Boosting Regressor^98^ with default hyperparameters in Scikit-learn, which was trained on half of the trials in each trial type condition and was tested on the other half of the trials for each animal. Fig. 4D compares the neural prediction of the latency to the probe-evoked swim from a randomly selected pulse with the ground truth in testing data for each trial condition across animals.

#### Statistical analysis

Data were evaluated for statistical significance using the following tests:

- Rank sum test (two-tailed): Fig. 1F, I, Fig. 3B
- Paired signed rank test: Fig. 1G, Fig. 5K-N,Q, Fig. S1A, Fig. S2A,B,D, Fig. S6A
- Spearman’s rank correlation: Fig. 2B1, Fig. 2D1, Fig. 2E1, Fig. 2F1, Fig. 4C, Fig. 4F
- Two-way ANOVA to evaluate fish identity contribution: Fig. 2B4, Fig. 2C4, Fig. 2D4, Fig. 2E4, Fig. 2F4, Fig. 4M,N
- Chi-square test: Fig. 5J
- Linear regression: Fig. 3E, H

